# Hippocampal-parietal cortex causal directed connectivity during human episodic memory formation: Replication across three experiments

**DOI:** 10.1101/2023.11.07.566056

**Authors:** Anup Das, Vinod Menon

## Abstract

Hippocampus-parietal cortex circuits are thought to play a crucial role in memory and attention, but their neural basis remains poorly understood. We employed intracranial EEG from 96 participants (51 females) to investigate the neurophysiological underpinning of these circuits across three memory tasks spanning verbal and spatial domains. We uncovered a consistent pattern of higher causal directed connectivity from the hippocampus to both lateral parietal cortex (supramarginal and angular gyrus) and medial parietal cortex (posterior cingulate cortex) in the delta-theta band during memory encoding and recall. This connectivity was independent of activation or suppression states in the hippocampus or parietal cortex. Crucially, directed connectivity from the supramarginal gyrus to the hippocampus was enhanced in participants with higher memory recall, highlighting its behavioral significance. Our findings align with the attention-to-memory model, which posits that attention directs cognitive resources toward pertinent information during memory formation. The robustness of these results was demonstrated through Bayesian replication analysis of the memory encoding and recall periods across the three tasks. Our study sheds light on the neural basis of casual signaling within hippocampus-parietal circuits, broadening our understanding of their critical roles in human cognition.

## Introduction

Episodic memory, the recollection of specific events and experiences, is central to human cognition (Tulving, 2002). The formation of episodic memories is thought to be supported by distributed brain circuits that integrate neural and cognitive processes involved in encoding and recall of specific events and experiences (Moscovitch, Cabeza, Winocur, & Nadel, 2016). While the hippocampus has been long recognized as a key brain structure for memory formation (Burgess, Maguire, & O’Keefe, 2002), a growing body of research is uncovering a key role for the parietal cortex in memory recall and recollection, expanding our understanding of the neurocognitive processes involved in episodic memory (Cabeza, Ciaramelli, Olson, & Moscovitch, 2008; Curtis, 2006; Husain & Nachev, 2007; Rolls, 2018, 2019; Vogt & Pandya, 1987; Wagner, Shannon, Kahn, & Buckner, 2005; Whitlock, Sutherland, Witter, Moser, & Moser, 2008). However, the electrophysiological correlates of their dynamic circuit interactions and specific spectral-temporal characteristics of how hippocampus-parietal circuits operate during human memory formation remain poorly understood.

Although previous studies predominantly emphasized the significance of hippocampus-prefrontal circuits in episodic memory (Dickerson & Eichenbaum, 2010; Eichenbaum, 2017; Moscovitch et al., 2016), there is growing evidence that hippocampus-parietal circuits also play a pivotal role in episodic memory (Simons, Ritchey, & Fernyhough, 2022). Despite its importance, this circuitry has received comparatively less attention in the scientific literature, leaving a significant gap in our understanding of its functional relevance in memory encoding and retrieval. Our study aims to address this gap by investigating the electrophysiological dynamics between the hippocampus and the parietal cortex, thereby contributing to a more comprehensive understanding of the neural mechanisms underlying episodic memory in the human brain.

Here, we leverage a large sample of intracranial EEG (iEEG) recordings, from 96 participants and three separate memory experiments, and investigate the temporal dynamics of signaling between the hippocampus and multiple distinct anatomical subdivisions of the parietal cortex. Spanning memory encoding and recall across both verbal and spatial domains, our study aims to unveil the causal dynamics of hippocampal-parietal interactions, enriching our understanding of episodic memory’s neural architecture and bridging important gaps in our current knowledge.

Extensive research in rodents and non-human primates has laid the groundwork for understanding the role of the hippocampus and parietal cortex in spatial memory. Single-neuron studies in these models have identified the parietal cortex as a key neural substrate for spatial memory tasks (Andersen, Essick, & Siegel, 1985; Chen, Lin, Green, Barnes, & McNaughton, 1994; Crowe, Chafee, Averbeck, & Georgopoulos, 2004; McNaughton et al., 1994; Nitz, 2006). Further, lesion studies have not only reinforced the importance of the parietal cortex in memory tasks (Barrow & Latto, 1996; Goodrich-Hunsaker, Hunsaker, & Kesner, 2005; Rogers & Kesner, 2006), but have also highlighted the crucial connectivity between the hippocampus and the parietal cortex for successful memory retrieval (Rogers & Kesner, 2007). These interactions are likely subserved by direct anatomical pathways, as anterograde and retrograde tracing studies in non-human primates have uncovered bidirectional projections between the hippocampus and parietal cortex (Clower, West, Lynch, & Strick, 2001; Insausti & Muñoz, 2001; Rockland & Van Hoesen, 1999), reinforcing their interconnected relationship. However, a translational gap exists, as the electrophysiological correlates of hippocampus-parietal circuit function have not been fully explored in human studies, a limitation our research aims to address.

In human studies, functional magnetic resonance imaging (fMRI) has consistently shown that both the hippocampus and the lateral parietal cortex are engaged during memory encoding and recall, as evidenced by a multitude of studies (Auger & Maguire, 2013; Benoit & Schacter, 2015; Brodt et al., 2018; Brodt et al., 2016; Buckner et al., 1998; Ciaramelli, Burianová, Vallesi, Cabeza, & Moscovitch, 2020; Gurd et al., 2002; Jonker, Dimsdale-Zucker, Ritchey, Clarke, & Ranganath, 2018; Konishi, Wheeler, Donaldson, & Buckner, 2000; Korkki, Richter, & Simons, 2021; Leech & Sharp, 2014; Miyamoto et al., 2013; Spreng, Stevens, Chamberlain, Gilmore, & Schacter, 2010; Summerfield, Hassabis, & Maguire, 2010; Vincent et al., 2006). Event-related potentials using scalp EEG recordings have illuminated the temporal aspects of brain activity, highlighting the role of the parietal cortex in successful memory retrieval (Paller, Kutas, & McIsaac, 1995; Paller, McCarthy, Roessler, Allison, & Wood, 1992; Paller, McCarthy, & Wood, 1988; Rugg et al., 1998; Wilding & Rugg, 1996). Further substantiating this line of inquiry, magnetoencephalography studies have suggested dissociable roles for delta, theta, and beta rhythms within the parietal cortex during the recall of episodic memory (Osipova et al., 2006; Seibert, Hagler, & Brewer, 2011).

Building on this, human iEEG depth recordings have confirmed the lateral parietal cortex’s role in episodic memory encoding and recall, as reflected in enhanced high-gamma frequency band activity (Gonzalez et al., 2015; Rubinstein et al., 2021; Tan, Rugg, & Lega, 2020). These studies also reveal a theta frequency band correlation between hippocampus and parietal cortex in tasks involving virtual spatial navigation (Ekstrom et al., 2005). Complementing these findings, transcranial stimulation studies have illustrated enhanced information flow between the parietal cortex and the hippocampus, resulting in improved memory performance (Crossman, Bartl, Soerum, & Sandrini, 2019; J. X. Wang et al., 2014). However, the precise nature of bidirectional interactions between the hippocampus and lateral parietal cortex, along with their frequency specificity, remains largely unexplored. Critically, the question of whether hippocampus-parietal circuity operate similarly in the verbal and spatial domains has not been addressed. This leaves a critical gap in our understanding of how the hippocampus interacts with the various anatomical subdivisions of the parietal cortex to support human episodic memory across different cognitive contexts.

The human parietal cortex is marked by a unique heterogeneity, featuring subdivisions that have undergone significant evolutionary expansion in comparison to rodents and non-human primates (Goldring & Krubitzer, 2020). Both the lateral and medial parietal cortex have undergone significant expansion. On the lateral surface, the supramarginal gyrus is engaged in a wide range of tasks requiring spatial attention. In contrast, the angular gyrus is crucial for the retrieval of vivid details and schema-based information (Buckner, Andrews-Hanna, & Schacter, 2008; Corbetta, 1998; Rugg & Vilberg, 2013). Notably, neither rodent nor non-human primate brains include an angular gyrus subdivision leaving unclear the dynamic interactive roles these lateral parietal subdivisions play in episodic memory. Furthermore, the angular gyrus, together with the posterior cingulate cortex (PCC)/precuneus in the medial parietal cortex, serve as core nodes of the default mode network. This network is typically deactivated during tasks that demand external attention (Bressler & Menon, 2010; Garrison et al., 2013; Greicius, Krasnow, Reiss, & Menon, 2003; Raichle et al., 2001; Zhang & Li, 2012), further suggesting a demarcation of their roles in episodic memory. The anatomical and functional complexity of the human parietal cortex necessitates a more precise examination of its role in hippocampal-parietal circuitry for memory formation (Sestieri, Shulman, & Corbetta, 2017). Our study aims to fill this gap by investigating the electrophysiological dynamics between the hippocampus and key subdivisions of the parietal cortex, both in its lateral and medial aspects. This focus allows for a more comprehensive understanding of hippocampal-parietal circuits, addressing a critical, yet overlooked area in the existing literature.

We leveraged iEEG data from the UPENN-RAM consortium (Solomon et al., 2019) to address critical gaps in our understanding of the electrophysiological mechanisms underpinning hippocampal-parietal cortex interactions in memory formation. This dataset includes depth recordings from an extensive cohort of individuals, allowing us to rigorously investigate the directionality and frequency specificity of connectivity between the hippocampus and multiple subdivisions of the lateral and medial parietal cortex. We employed three distinct episodic memory experiments: (i) a verbal free recall (VFR) task, in which participants were presented with a sequence of words for subsequent verbal recall, (ii) a categorized verbal free recall (CATVFR) task, which similarly involved encoding and recalling a sequence of words, but with the added demands of categorical organization, and (iii) a spatial memory (SM) task, wherein participants were required to memorize the spatial location of the objects for subsequent recall. These comprehensive datasets afforded a unique opportunity to probe hippocampal-parietal cortex interactions across memory encoding and recall periods and multiple cognitive domains.

The first objective of our study was to delineate the directed, causal connectivity between the hippocampus and multiple subdivisions of the lateral and medial parietal cortex during verbal and spatial memory tasks. To accomplish this, we employed phase transfer entropy (PTE) (Hillebrand et al., 2016; Lobier, Siebenhühner, Palva, & Matias, 2014; M. Y. Wang et al., 2017). PTE provides a robust quantitative measure for characterizing directional connectivity, capturing linear as well as nonlinear dynamics in iEEG data (Hillebrand et al., 2016; Lobier et al., 2014; Menon et al., 1996).

We examined three main hypotheses to guide our analysis. First, based on the ‘top-down attention’ hypothesis related to the supramarginal gyrus in memory encoding (Cabeza, 2008; Cabeza, Ciaramelli, & Moscovitch, 2012; Cabeza et al., 2011; Ciaramelli, Grady, & Moscovitch, 2008; Daselaar et al., 2009; Hutchinson, Uncapher, & Wagner, 2009; Uncapher & Wagner, 2009), we predicted that causal influences from the supramarginal gyrus to the hippocampus would be more pronounced compared to the reverse influence during encoding. Second, based on the bottom-up attention hypothesis of the angular gyrus in memory recall (Cabeza et al., 2008), we hypothesized that causal influences from hippocampus to angular gyrus would be stronger compared to the reverse direction. Third, we extended our hypotheses to the interactions between the hippocampus and the PCC/precuneus. We posited that the hippocampus would exert stronger influence on the PCC/precuneus during memory recall, mirroring its dynamics with the angular gyrus. This hypothesis is grounded in the fact that both the PCC/precuneus and the angular gyrus are integral nodes of the DMN, a brain network known to be involved in episodic memory recall and internally-driven cognition (Menon, 2023).

Our second objective was to investigate the frequency-specificity of causal interactions between the hippocampus and parietal cortex. Prior research across various species, including rodents, non-human primates, and humans, have emphasized the prominent functional roles of distinct frequency bands. Specifically, the delta-theta rhythm (0.5-8 Hz) has been identified as crucial in hippocampal functions (Ekstrom & Watrous, 2014; Neuner et al., 2014; Watrous, Tandon, Conner, Pieters, & Ekstrom, 2013), while beta-band rhythms (12-30 Hz) have been highlighted in lateral parietal cortex operations (Brovelli et al., 2004; Engel & Fries, 2010; Spitzer & Haegens, 2017; Stanley, Roy, Aoi, Kopell, & Miller, 2018). It has been suggested that delta-theta oscillations may facilitate synchronization between hippocampus and parietal cortex (Ekstrom & Watrous, 2014), while beta band oscillations may serve to coordinate neuronal cortico-cortical signaling (Engel & Fries, 2010; Negrón-Oyarzo et al., 2018; Spitzer & Haegens, 2017). However, the frequency-specificity of directed causal interactions between the hippocampus and parietal cortex during verbal and spatial memory remains an open question. In previous research on hippocampal-prefrontal cortex circuits, information theoretic analysis revealed greater causal information flow from hippocampus to the prefrontal cortex in the delta-theta frequency band (0.5-8 Hz); in contrast, prefrontal cortex to hippocampus causal information flow were stronger in the beta band (12-30 Hz) (Das & Menon, 2021, 2022b). Building on this work, we examined whether a similar profile of directional interactions would hold in the case of hippocampal-parietal cortex circuits. We hypothesized that the hippocampus would exert a stronger causal influence on both the lateral and medial parietal cortex in the delta-theta band. Conversely, we posited that the lateral parietal cortex would have a stronger “top-down” causal influence on the hippocampus in the beta band. Additionally, we also examined directed connectivity between the hippocampus and the PCC/precuneus in both frequency bands.

Our third objective was to probe regional profiles of neural response in the hippocampus and the three specific parietal cortex subdivisions. High-gamma band activity reflects localized, task-related neural processing, and is associated with synchronized activity of local neural populations (Crone, Boatman, Gordon, & Hao, 2001; Daitch & Parvizi, 2018; Edwards, Soltani, Deouell, Berger, & Knight, 2005; Lachaux et al., 2005; Ray, Crone, Niebur, Franaszczuk, & Hsiao, 2008; Sederberg, Schulze-Bonhage, Madsen, Bromfield, McCarthy, et al., 2007; Tallon-Baudry, Bertrand, Hénaff, Isnard, & Fischer, 2005). Moreover, increases in high-gamma power correlate with elevated neuronal spiking and synaptic activity, making it a robust marker for task-specific computations within local neuronal circuits (Ray et al., 2008). We determined whether high-gamma band power in these parietal subdivisions exhibits context-dependent variations across two critical periods of memory formation—encoding and recall. Given that memory encoding is largely influenced by external stimuli, we anticipated distinct neural response patterns in this period compared to memory recall, which is mainly internally driven (Andrews-Hanna, 2012; Buckner et al., 2008). Specifically, we hypothesized that high-gamma power in the angular gyrus and the PCC/precuneus, regions associated with the DMN, would be reduced during encoding, reflecting the externally driven nature of the task. Conversely, we expected an increase in high-gamma power during recall, which requires more internally oriented cognition. Furthermore, we expected the supramarginal gyrus, which is more engaged in tasks requiring adaptive external response, to show differential modulations in high-gamma power during both encoding and recall periods. These analyses aimed to provide complementary information on the differential roles played by these brain regions in the orchestration of episodic memory formation.

Our final objective was to assess the robustness and replicability of our findings across three distinct episodic memory experiments, encompassing the verbal and spatial domains. Replicability poses a significant challenge in the field of neuroscience, and this issue is particularly acute in invasive iEEG studies, where difficulty of acquiring data, small sample sizes, and lack of data sharing often hinder the generalizability of findings. To address this ‘reproducibility crisis’, we employed Bayesian analysis which provides a robust quantitative approach that allows for the explicit quantification of evidence for or against a given hypothesis (Ly, Etz, Marsman, & Wagenmakers, 2019; Verhagen & Wagenmakers, 2014). Specifically, we used Bayes factors (BFs) to evaluate the strength and consistency of our findings across verbal and spatial memory tasks. BF provides a powerful metric for comparing the likelihood of the observed data under different hypotheses, offering a rigorous measure of the evidence supporting the replication of effects. This analysis allowed us to provide a comprehensive assessment of the generalizability of our findings.

Our study offers new insights into the neurophysiological mechanisms that govern cognitive processes, specifically highlighting the role of hippocampal-parietal cortex interactions in both encoding and recall of verbal and spatial information. Crucially, the robustness and generalizability of our findings were confirmed through their replication across three distinct experiments spanning verbal and spatial domains, emphasizing the importance of hippocampal-parietal interactions in a broader context of human cognition.

## Materials and Methods

### UPENN-RAM iEEG recordings

iEEG recordings from 249 patients shared by Kahana and colleagues at the University of Pennsylvania (UPENN) (obtained from the UPENN-RAM public data release) were used for analysis (Jacobs et al., 2016). Patients with pharmaco-resistant epilepsy underwent surgery for removal of their seizure onset zones. iEEG recordings of these patients were downloaded from a UPENN-RAM consortium hosted data sharing archive (URL: http://memory.psych.upenn.edu/RAM). Prior to data collection, research protocols and ethical guidelines were approved by the Institutional Review Board at the participating hospitals and informed consent was obtained from the participants and guardians (Jacobs et al., 2016). Details of all the recordings sessions and data pre-processing procedures are described by Kahana and colleagues (Jacobs et al., 2016). Briefly, iEEG recordings were obtained using subdural grids and strips (contacts placed 10 mm apart) or depth electrodes (contacts spaced 5–10 mm apart) using recording systems at each clinical site. iEEG systems included DeltaMed XlTek (Natus), Grass Telefactor, and Nihon-Kohden EEG systems. Electrodes located in brain lesions or those which corresponded to seizure onset zones or had significant interictal spiking or had broken leads, were excluded from analysis.

Anatomical localization of electrode placement was accomplished by co-registering the postoperative computed CTs with the postoperative MRIs using FSL (FMRIB (Functional MRI of the Brain) Software Library), BET (Brain Extraction Tool), and FLIRT (FMRIB Linear Image Registration Tool) software packages. Preoperative MRIs were used when postoperative MRIs were not available. The resulting contact locations were mapped to MNI space using an indirect stereotactic technique and OsiriX Imaging Software DICOM viewer package. We used the Brainnetome atlas (Fan et al., 2016) to demarcate the left hemisphere subdivisions of the parietal cortex (supramarginal gyrus (SMG), angular gyrus (AG), and posterior cingulate cortex (PCC)/precuneus) (Cabeza et al., 2008; Hutchinson et al., 2009) and the hippocampus. We first identified electrode pairs in which there were at least four patients with electrodes implanted in each pair of brain regions of interest encompassing SMG, AG, PCC/precuneus, and hippocampus (**Tables 2, 3**). The lack of sufficient number of participants and electrode pairs precluded analyses of two other important subdivisions of the lateral parietal cortex namely, the superior parietal lobule and the intraparietal sulcus. Out of 249 individuals, data from 96 individuals (aged from 18 to 64, mean age 38.6 ± 12.3, 51 females) were used for subsequent analysis based on electrode placement in supramarginal gyrus, angular gyrus, PCC/precuneus, and hippocampus. Gender differences were not analyzed in this study due to lack of sufficient male participants for electrodes pairs for hippocampus-SMG, hippocampus-AG, and hippocampus-PCC/precuneus interactions (**Table 2c**).

Original sampling rates of iEEG signals were 500 Hz, 1000 Hz, 1024 Hz, and 1600 Hz. Hence, iEEG signals were downsampled to 500 Hz, if the original sampling rate was higher, for all subsequent analysis. The two major concerns when analyzing interactions between closely spaced intracranial electrodes are volume conduction and confounding interactions with the reference electrode (Burke et al., 2013). Hence bipolar referencing was used to eliminate confounding artifacts and improve the signal-to-noise ratio of the neural signals, consistent with previous studies using UPENN-RAM iEEG data (Burke et al., 2013; Ezzyat et al., 2018). Signals recorded at individual electrodes were converted to a bipolar montage by computing the difference in signal between adjacent electrode pairs on each strip, grid, and depth electrode and the resulting bipolar signals were treated as new “virtual” electrodes originating from the midpoint between each contact pair, identical to procedures in previous studies using UPENN-RAM data (Solomon et al., 2019). Line noise (60 Hz) and its harmonics were removed from the bipolar signals using a fourth order two-way zero phase lag Butterworth filter and finally each bipolar signal was Z-normalized by removing mean and scaling by the standard deviation. For filtering, we used a fourth order two-way zero phase lag Butterworth filter throughout the analysis.

### iEEG verbal free recall and spatial memory tasks

#### (a) Verbal free recall (VFR) task

Patients performed multiple trials of a “free recall” experiment, where they were presented with a list of words and subsequently asked to recall as many as possible from the original list (**Figure 1a**) (Solomon et al., 2017; Solomon et al., 2019). The task consisted of three periods: encoding, delay, and recall. During encoding, a list of 12 words was visually presented for ∼30 sec. Words were selected at random, without replacement, from a pool of high frequency English nouns (http://memory.psych.upenn.edu/Word_Pools). Each word was presented for a duration of 1600 msec, followed by an inter-stimulus interval of 800 to 1200 msec. After a 20 sec post-encoding delay, participants were instructed to recall as many words as possible during the 30 sec recall period. Average recall accuracy across patients was 24.8% ± 9.7%, similar to prior studies of verbal episodic memory retrieval in neurosurgical patients (Burke et al., 2014). The mismatch in the number trials therefore made it difficult to directly compare causal signaling measures between successfully versus unsuccessfully recalled words. From the point of view of probing behaviorally effective memory encoding, our focus was therefore on successful encoding and recall consistent with most prior studies (Long, Burke, & Kahana, 2014; Watrous et al., 2013). We analyzed iEEG epochs from the encoding and recall periods of the free recall task. For the encoding periods, 1600 msec iEEG recordings immediately following the stimulus presentation were analyzed (Solomon et al., 2019). For the recall periods, iEEG recordings 1600 msec prior to the vocal onset of each word were analyzed (Solomon et al., 2019). Recall epochs which overlapped with the prior recall epoch were excluded from analysis. Data from each trial was analyzed separately and specific measures were averaged across trials.

**Figure 1.**
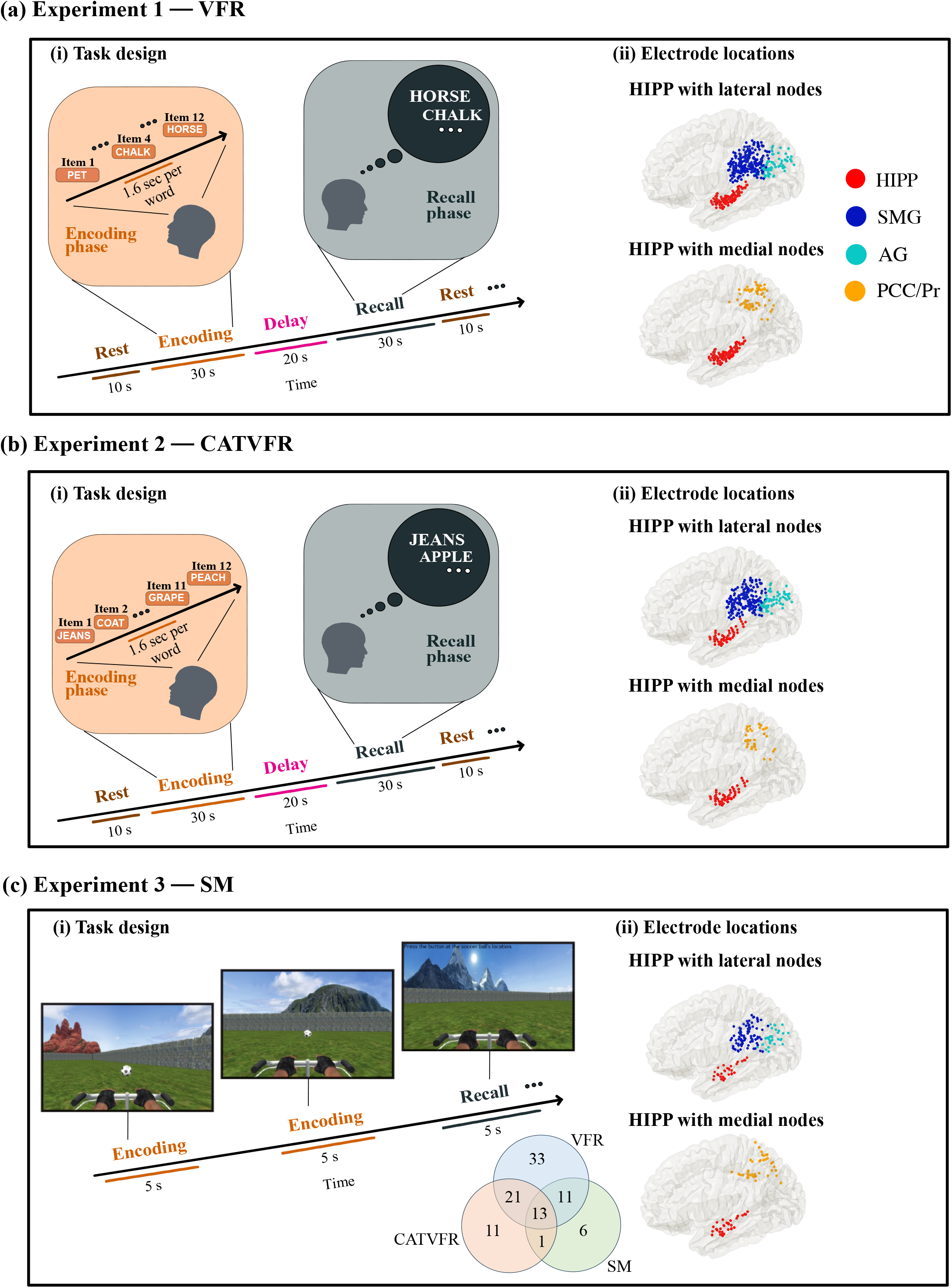
Task design of the encoding and recall periods of the memory experiments, and iEEG recording sites in the HIPP, with the lateral and medial nodes of the parietal cortex. **(a) Experiment 1, Verbal free recall (VFR): (i)** Task design of memory encoding and recall periods of the verbal free recall experiment (see **Methods** for details). Participants were first presented with a list of words in the encoding block and asked to recall as many as possible from the original list after a short delay. **(ii)** Electrode locations for HIPP with lateral nodes (top panel) and HIPP with medial nodes (bottom panel), in the verbal free recall experiment. **(b) Experiment 2, Categorized verbal free recall (CATVFR): (i)** Task design of memory encoding and recall periods of the categorized verbal free recall experiment (see **Methods** for details). Participants were presented with a list of words with consecutive pairs of words from a specific category (for example, JEANS-COAT, GRAPE-PEACH, etc.) in the encoding block and subsequently asked to recall as many as possible from the original list after a short delay. **(ii)** Electrode locations for HIPP with lateral nodes (top panel) and HIPP with medial nodes (bottom panel), in the categorized verbal free recall experiment. **(c) Experiment 3, Spatial memory (SM): (i)** Task design of memory encoding and recall periods of the spatial memory experiment (see **Methods** for details). Participants were shown objects in various locations during the encoding period and asked to retrieve the location of the objects during the recall period. Modified from Jacobs et. al. (2018) with permission. **(ii)** Electrode locations for HIPP with lateral nodes (top panel) and HIPP with medial nodes (bottom panel), in the spatial memory experiment. Venn diagram shows overlap of the number of subjects across the three experiments. HIPP: hippocampus, SMG: supramarginal gyrus, AG: angular gyrus, PCC: posterior cingulate cortex, Pr: precuneus.

#### (b) Categorized verbal free recall (CATVFR) task

This task was very similar to the verbal free recall task. Here, patients performed multiple trials of a “categorized free recall” experiment, where they were presented with a list of words with consecutive pairs of words from a specific category (for example, JEANS-COAT, GRAPE-PEACH, etc.) and subsequently asked to recall as many as possible from the original list (**Figure 1b**) (Qasim, Mohan, Stein, & Jacobs, 2023). Similar to the uncategorized verbal free recall task, this task also consisted of three periods: encoding, delay, and recall. During encoding, a list of 12 words was visually presented for ∼30 sec. Semantic categories were chosen using Amazon Mechanical Turk. Pairs of words from the same semantic category were never presented consecutively. Each word was presented for a duration of 1600 msec, followed by an inter-stimulus interval of 750 to 1000 msec. After a 20 sec post-encoding delay, participants were instructed to recall as many words as possible during the 30 sec recall period. Average recall accuracy across patients was 27.0% ± 11.9%. Similar to the uncategorized verbal free recall task, we focused on trials corresponding to successful encoding and recall. Analysis of iEEG epochs from the encoding and recall periods of the categorized free recall task was same as the uncategorized verbal free recall task.

#### (c) Spatial memory (SM) task

Participants performed multiple trials of a spatial memory task in a virtual navigation paradigm (Goyal et al., 2018; Jacobs et al., 2016; Lee et al., 2018) similar to the Morris water maze (Morris, 1984). The environment was rectangular (1.8:1 aspect ratio), and was surrounded by a continuous boundary (**Figure 1c**). There were four distal visual cues (landmarks), one centered on each side of the rectangle, to aid with orienting. Each trial (96 trials per session, 1–3 sessions per subject) started with two 5 sec *encoding periods*, during which subjects were driven to an object from a random starting location. At the beginning of an encoding period, the object appeared and, over the course of 5 sec, the subject was automatically driven directly toward it. The 5 sec period consisted of three intervals: first, the subject was rotated toward the object (1 sec), second, the subject was driven toward the object (3 sec), and, finally, the subject paused while at the object location (1 sec). After a 5 sec delay with a blank screen, the same process was repeated from a different starting location. After both encoding periods for each item, there was a 5 sec pause followed by the *recall period*. The subject was placed in the environment at a random starting location with the object hidden and then asked to freely navigate using a joystick to the location where they thought the object was located. When they reached their chosen location, they pressed a button to record their response. They then received feedback on their performance via an overhead view of the environment showing the actual and reported object locations. Average recall accuracy across patients was 48.6% ± 4.2%.

Similar to the verbal free recall task, only trials corresponding to successful memory encoding and recall are considered in our analysis. We analyzed 5 sec iEEG epochs from the encoding and recall periods of the task. Data from each trial was analyzed separately and specific measures were averaged across trials, similar to the verbal free recall task.

### iEEG analysis of high-gamma power

We first filtered the signals in the high-gamma (80-160 Hz) frequency band (Raccah, Daitch, Kucyi, & Parvizi, 2018) and then calculated the square of the filtered signals as the power of the signals (Kwon et al., 2021). Signals were then smoothed using 0.2s windows with 90% overlap (Kwon et al., 2021) and normalized with respect to 0.2s pre-stimulus periods.

### iEEG analysis of phase transfer entropy (PTE) and causal dynamics

Phase transfer entropy (PTE) is a nonlinear measure of the directionality of connectivity between time-series and can be applied as a measure of causality to nonstationary time-series (Lobier et al., 2014). The PTE measure is in contrast to the Granger causality measure which can be applied only to stationary time-series (Barnett & Seth, 2014). We first carried out a stationarity test of the iEEG recordings (unit root test for stationarity (Barnett & Seth, 2014)) and found that the spectral radius of the autoregressive model is very close to one, indicating that the iEEG time-series is nonstationary. This precluded the applicability of the Granger causality analysis in our study.

Given two time-series {*x_i_* } and {*y_i_* }, where *i* = 1,2,…, *M*, instantaneous phases were first extracted using the Hilbert transform. Let 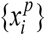 and 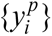, where *i* = 1,2,…, *M*, denote the corresponding phase time-series. If the uncertainty of the target signal 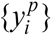 at delay *τ* is quantified using Shannon entropy, then the PTE from driver signal 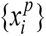 to target signal 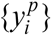 can be given by

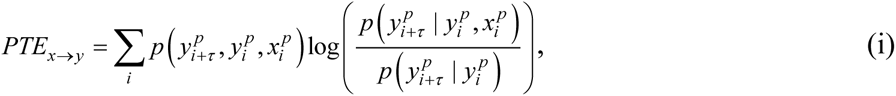

where the probabilities can be calculated by building histograms of occurrences of singles, pairs, or triplets of instantaneous phase estimates from the phase time-series (Hillebrand et al., 2016). For our analysis, the number of bins in the histograms was set as 3.49×*STD*×*M* ^−1/3^ and delay *τ* was set as 2*M* / *M*_±_, where *STD* is average standard deviation of the phase time-series 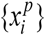 and 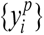 and *M*_±_ is the number of times the phase changes sign across time and channels (Hillebrand et al., 2016). PTE has been shown to be robust against the choice of the delay *τ* and the number of bins for forming the histograms (Hillebrand et al., 2016).

### Statistical analysis

Statistical analysis was conducted using mixed effects analysis with the lmerTest package (Kuznetsova, Brockhoff, & Christensen, 2017) implemented in R software (version 4.0.2, R Foundation for Statistical Computing). Because PTE data were not normally distributed, we used BestNormalize (Peterson & Cavanaugh, 2018) which contains a suite of transformation-estimating functions that can be used to optimally normalize data. The resulting normally distributed data were subjected to mixed effects analysis with the following model: *PTE ∼ Condition + (1|Subject)*, where *Condition* models the fixed effects (condition differences) and (1|*Subject*) models the random repeated measurements within the same participant. Before running the mixed-effects model, PTE was first averaged across trials for each channel pair. We modeled brain region pairs (**Figure 3, 5**, for example, HIPP → SMG versus SMG → HIPP), performance (**Figure 6**, higher versus lower performing individuals), or individual brain regions (**Figure 7**, for example, HIPP versus SMG) as fixed effects. Analysis of variance (ANOVA) was used to test the significance of findings with FDR-corrections for multiple comparisons across task conditions, frequencies, and brain regions (*p*<0.05). Similar mixed effects statistical analysis procedures were used for comparison of high-gamma power across task conditions, where the mixed effects analysis was run on each of the 0.2s windows.

Finally, we conducted surrogate analysis to test the significance of the estimated PTE values (Hillebrand et al., 2016). The estimated phases from the Hilbert transform for electrodes from a given pair of brain areas were time-shuffled so that the predictability of one time-series from another is destroyed, and PTE analysis was repeated on this shuffled data to build a distribution of surrogate PTE values against which the observed PTE was tested (*p*<0.05).

### Bayesian replication analysis

We used replication Bayes factor (Ly et al., 2019; Verhagen & Wagenmakers, 2014) analysis to estimate the degree of replicability for the direction of causal influence for each frequency and task condition and across task domains. Analysis was implemented in R software using the BayesFactor package (Rouder, Speckman, Sun, Morey, & Iverson, 2009). Because PTE data were not normally distributed, as previously, we used BestNormalize (Peterson & Cavanaugh, 2018) to optimally normalize data. We calculated the replication Bayes factor for pairwise task domains. We compared the Bayes factor of the joint model *PTE(task1+task2) ∼ Condition + (1|Subject)* with the Bayes factor (BF) of individual model as *PTE(task1) ∼ Condition + (1|Subject)*, where *task1* denotes the verbal free recall (original) task and *task2* denotes the categorized verbal free recall, or spatial memory (replication) conditions. We calculated the ratio *BF(task1+task2)/BF(task1)*, which was used to quantify the degree of replicability. We determined whether the degree of replicability was higher than 3 as Bayes factor of at least 3 indicates evidence for replicability (Jeffreys, 1998). A Bayes factor of at least 100 is considered as “*decisive*” for degree of replication (Jeffreys, 1998).

## Results

### Causal influences from the hippocampus to parietal cortex in delta-theta band during memory encoding

Based on previous iEEG studies which have reported significant delta-theta frequency (0.5-8 Hz) band activity in the hippocampus during recall of information from recently encoded memories (Ekstrom & Watrous, 2014; Watrous et al., 2013) and based on our recent findings where we found higher directed causal influence from the hippocampus to the prefrontal cortex than the reverse, in delta-theta frequency band during both encoding and recall, and across verbal and spatial episodic memory domains (Das & Menon, 2021, 2022b), we first investigated the dynamic causal influences of the hippocampus on SMG, AG, and PCC/precuneus and vice-versa in the low frequency delta-theta band. We computed PTE from SMG, AG, PCC/precuneus to the hippocampus and, in the reverse direction, during episodic memory encoding, and recall in the delta-theta band.

#### VFR task

Participants were presented with a sequence of words and asked to remember them for subsequent recall (**Methods**, **Tables 1, 2a, 3a, Figure 1a**). Briefly, the task consisted of three periods: encoding, delay, and recall (Solomon et al., 2019). During encoding, a list of 12 words was visually presented for ∼30 sec. Each word was presented for a duration of 1600 msec, followed by an inter-stimulus interval of 800 to 1200 msec. After a 20 sec post-encoding delay, participants were instructed to recall as many words as possible during the 30 sec recall period.

**Table 1.** Participant demographic information (total 96 participants).

| **Participant ID** | **Gender** | **Age** |
| --- | --- | --- |
| 002 | F | 49 |
| 003 | F | 39 |
| 006 | F | 20 |
| 008 | F | 55 |
| 010 | F | 30 |
| 014 | F | 47 |
| 015 | F | 54 |
| 018 | M | 47 |
| 021 | M | 34 |
| 024 | F | 36 |
| 030 | M | 23 |
| 032 | F | 19 |
| 033 | F | 31 |
| 034 | F | 29 |
| 035 | F | 45 |
| 036 | M | 49 |
| 044 | M | 58 |
| 049 | F | 52 |
| 050 | M | 20 |
| 051 | F | 24 |
| 052 | F | 19 |
| 054 | M | 23 |
| 059 | F | 44 |
| 063 | M | 23 |
| 065 | F | 34 |
| 066 | M | 39 |
| 067 | F | 45 |
| 069 | M | 26 |
| 070 | F | 40 |
| 073 | M | 60 |
| 074 | M | 24 |
| 075 | M | 50 |
| 076 | M | 29 |
| 080 | F | 43 |
| 081 | F | 33 |
| 083 | F | 49 |
| 084 | M | 25 |
| 089 | M | 36 |
| 092 | M | 44 |
| 094 | M | 47 |
| 101 | F | 26 |
| 102 | M | 34 |
| 111 | M | 20 |
| 114 | F | 31 |
| 119 | F | 26 |
| 120 | F | 33 |
| 122 | F | 48 |
| 124 | F | 40 |
| 125 | F | 44 |
| 130 | M | 57 |
| 134 | M | 64 |
| 135 | M | 47 |
| 137 | F | 21 |
| 138 | M | 41 |
| 144 | M | 53 |
| 147 | M | 47 |
| 151 | M | 36 |
| 153 | M | 38 |
| 154 | F | 36 |
| 158 | F | 45 |
| 161 | F | 53 |
| 163 | M | 45 |
| 164 | M | 37 |
| 168 | M | 24 |
| 174 | M | 29 |
| 175 | M | 34 |
| 177 | F | 23 |
| 180 | F | 21 |
| 187 | F | 51 |
| 190 | F | 57 |
| 193 | M | 37 |
| 195 | M | 44 |
| 196 | M | 18 |
| 203 | F | 36 |
| 204 | F | 25 |
| 207 | F | 39 |
| 221 | M | 57 |
| 223 | F | 42 |
| 227 | M | 32 |
| 228 | F | 58 |
| 230 | F | 56 |
| 232 | M | 27 |
| 234 | M | 25 |
| 236 | F | 51 |
| 239 | M | 27 |
| 240 | F | 37 |
| 245 | M | 30 |
| 247 | F | 61 |
| 260 | F | 57 |
| 279 | F | 57 |
| 286 | F | 57 |
| 292 | F | 39 |
| 297 | M | 24 |
| 298 | F | 24 |
| 299 | M | 43 |
| 302 | M | 48 |

**Table 2a.** Number of electrode pairs used in phase transfer entropy (PTE) analysis in the verbal free recall task. HIPP: hippocampus; SMG: supramarginal gyrus; AG: angular gyrus; PCC: posterior cingulate cortex, Pr: precuneus.

| Network pairs | Number of electrode pairs (n) | Number of participants | Participant IDs (Gender/Age) |
| --- | --- | --- | --- |
| HIPP-SMG | 184 | 23 | 003 (F/39), 010 (F/30), 035 (F/45), 049 (F/52), 054 (M/23), 065 (F/34), 066 (M/39), 080 (F/43), 111 (M/20), 114 (F/31), 120 (F/33), 134 (M/64), 135 (M/47), 138 (M/41), 151 (M/36), 161 (F/53), 164 (M/37), 174 (M/29), 195 (M/44), 236 (F/51), 240 (F/37), 297 (M/24), 299 (M/43) |
| HIPP-AG | 72 | 13 | 010 (F/30), 032 (F/19), 033 (F/31), 111 (M/20), 120 (F/33), 134 (M/64), 154 (F/36), 161 (F/53), 174 (M/29), 195 (M/44), 203 (F/36), 230 (F/56), 240 (F/37) |
| HIPP-PCC/Pr | 53 | 12 | 010 (F/30), 049 (F/52), 054 (M/23), 114 (F/31), 134 (M/64), 135 (M/47), 138 (M/41), 203 (F/36), 236 (F/51), 240 (F/37), 297 (M/24), 299 (M/43) |

**Table 3a.** Number of electrodes in each node used in high-gamma power analysis in the verbal free recall task. HIPP: hippocampus; SMG: supramarginal gyrus; AG: angular gyrus; PCC: posterior cingulate cortex, Pr: precuneus.

| Brain regions | Number of electrodes (n) | Number of participants | Participant IDs (Gender/Age) |
| --- | --- | --- | --- |
| HIPP | 108 | 40 | 002 (F/49), 003 (F/39), 010 (F/30), 032 (F/19), 033 (F/31), 035 (F/45), 049 (F/52), 054 (M/23), 063 (M/23), 065 (F/34), 066 (M/39), 080 (F/43), 089 (M/36), 111 (M/20), 114 (F/31), |
|  |  |  | 120 (F/33), 122 (F/48), 134 (M/64), 135 (M/47), 137 (F/21), 138 (M/41), 151 (M/36), 154 (F/36), 161 (F/53), 164 (M/37), 168 (M/24), 174 (M/29), 195 (M/44), 203 (F/36), 207 (F/39), 223 (F/42), 228 (F/58), 230 (F/56), 236 (F/51), 240 (F/37), 247 (F/61), 292 (F/39), 297 (M/24), 298 (F/24), 299 (M/43) |
| SMG | 230 | 54 | 003 (F/39), 010 (F/30), 015 (F/54), 030 (M/23), 035 (F/45), 036 (M/49), 044 (M/58), 049 (F/52), 050 (M/20), 054 (M/23), 059 (F/44), 065 (F/34), 066 (M/39), 067 (F/45), 069 (M/26), 070 (F/40), 074 (M/24), 075 (M/50), 080 (F/43), 084 (M/25), 094 (M/47), 102 (M/34), 111 (M/20), 114 (F/31), 120 (F/33), 125 (F/44), 130 (M/57), 134 (M/64), 135 (M/47), 138 (M/41), 147 (M/47), 151 (M/36), 153 (M/38), 158 (F/45), 161 (F/53), 163 (M/45), 164 (M/37), 174 (M/29), 177 (F/23), 193 (M/37), 195 (M/44), 204 (F/25), 232 (M/27), 234 (M/25), 236 (F/51), 240 (F/37), 260 (F/57), 286 (F/57), 297 (M/24), 299 (M/43) |
| AG | 72 | 29 | 010 (F/30), 032 (F/19), 033 (F/31), 034 (F/29), 036 (M/49), 050 (M/20), 059 (F/44), 069 (M/26), 070 (F/40), 081 (F/33), 083 (F/49), 084 (M/25), 094 (M/47), 101 (F/26), 111 (M/20), 120 (F/33), 134 (M/64), 147 (M/47), 154 (F/36), 158 (F/45), 161 (F/53), 174 (M/29), 187 (F/51), 195 (M/44), 196 (M/18), 203 (F/36), 230 (F/56), 240 (F/37), 260 (F/57) |
| PCC/Pr | 76 | 31 | 006 (F/20), 010 (F/30), 018 (M/47), 030 (M/23), 044 (M/58), 049 (F/52), 051 (F/24), 054 (M/23), 070 (F/40), 074 (M/24), 076 (M/29), 081 (F/33), 084 (M/25), 094 (M/47), 101 (F/26), 114 (F/31), 134 (M/64), 135 (M/47), 138 (M/41), 153 (M/38), 158 (F/45), 175 (M/34), 193 (M/37), 196 (M/18), 203 (F/36), 204 (F/25), 236 (F/51), |
|  |  |  | 240 (F/37), 286 (F/57), 297 (M/24),<br>299 (M/43) |

Surrogate data analysis revealed that causal influence from the hippocampus to the SMG, AG, and PCC/precuneus and the reverse were significantly higher than those expected by chance (*p*<0.05 in all cases) in the delta-theta frequency band, indicating bidirectional causal influence between the hippocampus and the parietal cortex in delta-theta band **(Figure 2a).** Next, we tested the hypothesis that the strength of causal influences from the hippocampus to parietal cortex would be higher than the reverse in the delta-theta band. This analysis revealed that the hippocampus had higher causal influences on SMG (*F*(1, 341) = 110.78, *p*<0.001, Cohen’s *d* = 1.14), AG (*F*(1, 130) = 69.65, *p*<0.001, Cohen’s *d* = 1.47), and PCC/precuneus (*F*(1, 93) = 72.97, *p*<0.001, Cohen’s *d* = 1.77) than the reverse, during memory encoding (**Figure 3a**).

**Figure 2.**
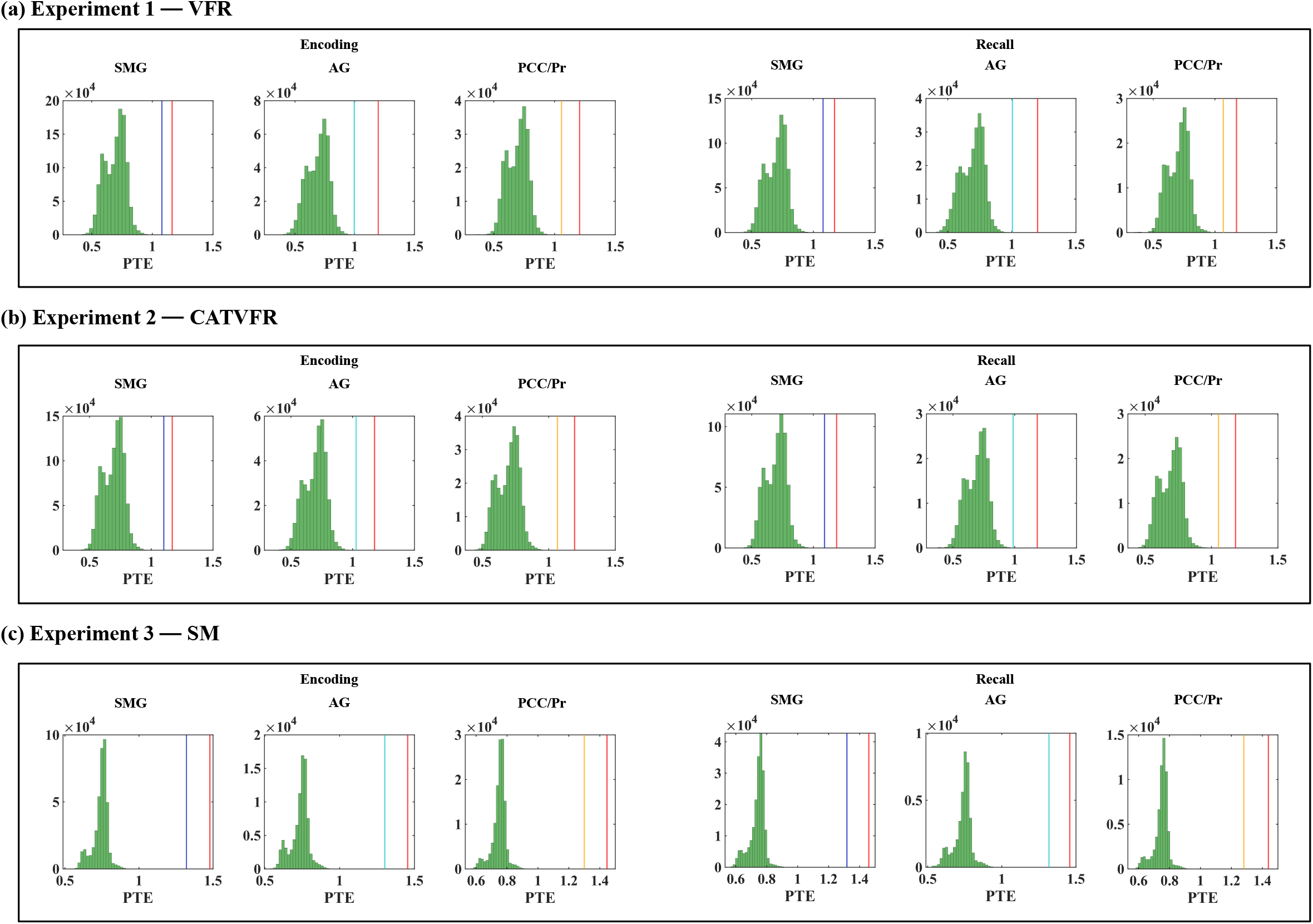
Surrogate data analysis results related to directed causal influence from the hippocampus to the parietal cortex (red) and the reverse (blue for supramarginal gyrus, cyan for angular gyrus, and orange for PCC/precuneus) in the delta-theta band, during (a) VFR, (b) CATVFR, and (c) SM experiments. Surrogate data analysis revealed that causal influence from the hippocampus to the SMG, AG, and PCC/precuneus and the reverse were significantly enhanced compared to those expected by chance during both encoding and recall periods, and across the three experiments (all *ps* < 0.05).

**Figure 3.**
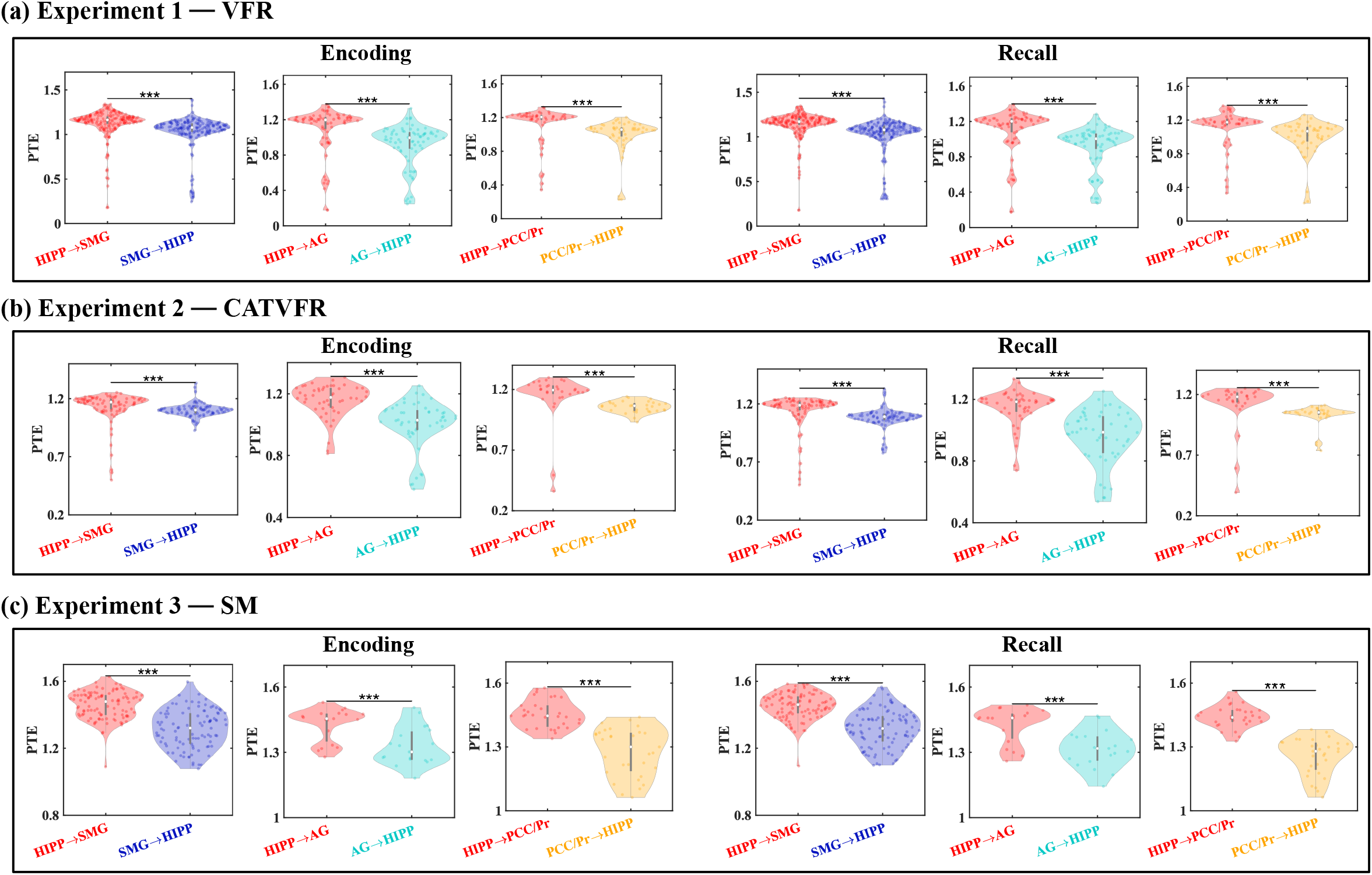
Directed connectivity between hippocampus and SMG, AG, and PCC/precuneus in delta-theta band (0.5-8 Hz). **(a) Experiment 1, VFR:** The hippocampus showed higher causal influences on the SMG (HIPP → SMG) (n=184), AG (HIPP → AG) (n=72), and PCC/precuneus (HIPP → PCC/Pr) (n=53) during both memory encoding and recall, compared to the reverse direction (SMG → HIPP, AG → HIPP, and PCC/Pr → HIPP respectively). **(b) Experiment 2, CATVFR:** The hippocampus showed higher causal directed influence on the SMG (HIPP → SMG) (n=87), AG (HIPP → AG) (n=50), and PCC/precuneus (HIPP → PCC/Pr) (n=31) during both memory encoding and recall, compared to the reverse direction (SMG → HIPP, AG → HIPP, and PCC/Pr → HIPP respectively). **(c) Experiment 3, SM:** The hippocampus also showed higher causal influences on the SMG (HIPP → SMG) (n=100), AG (HIPP → AG) (n=19), and PCC/precuneus (HIPP → PCC/Pr) (n=31) during both memory encoding and recall, compared to the reverse direction (SMG → HIPP, AG → HIPP, and PCC/Pr → HIPP respectively). *** *p* < 0.001 (Linear mixed effects ANOVA, FDR-corrected).

#### CATVFR task

We next examined directed causal influence from the hippocampus to the SMG, AG, and PCC/precuneus during memory encoding in the categorized verbal free recall task. In the CATVFR task, participants were presented with a list of words with consecutive pairs of words from a specific category (for example, JEANS-COAT, GRAPE-PEACH, etc.) and subsequently asked to recall as many as possible from the original list (**Methods**, **Tables 1, 2b, 3b**, **Figure 1b**) (Qasim et al., 2023). Similar to the uncategorized verbal free recall task, this task also consisted of three periods: encoding, delay, and recall. During encoding, a list of 12 words was visually presented for ∼30 sec. Each word was presented for a duration of 1600 msec, followed by an inter-stimulus interval of 750 to 1000 msec. After a 20 sec post-encoding delay, participants were instructed to recall as many words as possible during the 30 sec recall period.

**Table 2b.** Number of electrode pairs used in phase transfer entropy (PTE) analysis in the categorized verbal free recall task. HIPP: hippocampus; SMG: supramarginal gyrus; AG: angular gyrus; PCC: posterior cingulate cortex, Pr: precuneus.

| Network pairs | Number of electrode pairs (n) | Number of participants | Participant IDs (Gender/Age) |
| --- | --- | --- | --- |
| HIPP-SMG | 87 | 11 | 035 (F/45), 065 (F/34), 066 (M/39), 092 (M/44), 111 (M/20), 114 (F/31), 119 (F/26), 135 (M/47), 144 (M/53), 174 (M/29), 240 (F/37) |
| HIPP-AG | 50 | 7 | 021 (M/38), 032 (F/19), 111 (M/20), 119 (F/26), 174 (M/29), 230 (F/56), 240 (F/37) |
| HIPP-PCC/Pr | 31 | 4 | 114 (F/31), 135 (M/47), 240 (F/37), 245 (M/30) |

**Table 3b.** Number of electrodes in each node used in high-gamma power analysis in the categorized verbal free recall task. HIPP: hippocampus; SMG: supramarginal gyrus; AG: angular gyrus; PCC: posterior cingulate cortex, Pr: precuneus.

| Brain regions | Number of electrodes (n) | Number of participants | Participant IDs (Gender/Age) |
| --- | --- | --- | --- |
| HIPP | 55 | 22 | 021 (M/38), 032 (F/19), 035 (F/45), 065 (F/34), 066 (M/39), 089 (M/36), 092 (M/44), 111 (M/20), 114 (F/31), 119 (F/26), 135 (M/47), 144 (M/53), 174 (M/29), 180 (F/21), 207 (F/39), 228 (F/58), 230 (F/56), 236 (F/51), 239 (M/27), 240 (F/37), 245 (M/30), 247 (F/61) |
| SMG | 181 | 36 | 015 (F/54), 035 (F/45), 036 (M/49), 044 (M/58), 050 (M/20), 065 (F/34), 066 (M/39), 067 (F/45), 069 (M/26), 074 (M/24), 075 (M/50), 092 (M/44), 094 (M/47), 102 (M/34), 111 (M/20), 114 (F/31), 119 (F/26), 130 (M/57), 135 (M/47), 144 (M/53), 147 (M/47), 158 (F/45), 163 (M/45), 174 (M/29), 190 (F/57), 204 (F/25), 221 (M/57), 227 (M/32), 236 (F/51), 240 (F/37), 260 (F/57), 279 (F/57), 286 (F/57), 302 (M/48) |
| AG | 63 | 19 | 021 (M/38), 032 (F/19), 036 (M/49), 050 (M/20), 069 (M/26), 083 (F/49), 094 (M/47), 111 (M/20), 119 (F/26), 147 (M/47), 158 (F/45), 174 (M/29), 187 (F/51), 190 (F/57), 221 (M/57), 230 (F/56), 240 (F/37), 260 (F/57), 302 (M/48) |
| PCC/Pr | 33 | 12 | 044 (M/58), 074 (M/24), 094 (M/47), 114 (F/31), 135 (M/47), 158 (F/45), 204 (F/25), 227 (M/32), 236 (F/51), 240 (F/37), 245 (M/30), 286 (F/57) |

Surrogate data analysis revealed that causal influence from the hippocampus to the SMG, AG, and PCC/precuneus and the reverse were significantly higher than those expected by chance (*p*<0.05 in all cases) in the CATVFR task, indicating bidirectional causal influence between the hippocampus and the parietal cortex during this task **(Figure 2b).** We then tested the hypothesis that the strength of causal influences from the hippocampus to parietal cortex would be higher than the reverse. This analysis revealed that the hippocampus had higher causal influences on SMG (*F*(1, 158) = 26.67, *p*<0.001, Cohen’s *d* = 0.82), AG (*F*(1, 83) = 44.08, *p*<0.001, Cohen’s *d* = 1.45), and PCC/precuneus (*F*(1, 57) = 35.86, *p*<0.001, Cohen’s *d* = 1.59) than the reverse (**Figure 3b**).

#### SM task

We next examined dynamic causal influences of the hippocampus on the parietal cortex in the spatial memory task. In this task (**Methods**, **Tables 1, 2c, 3c**, **Figure 1c**), participants performed multiple trials of a spatial memory task in a virtual navigation paradigm (Goyal et al., 2018; Jacobs et al., 2016; Lee et al., 2018) similar to the Morris water maze (Morris, 1984).

**Table 2c.**
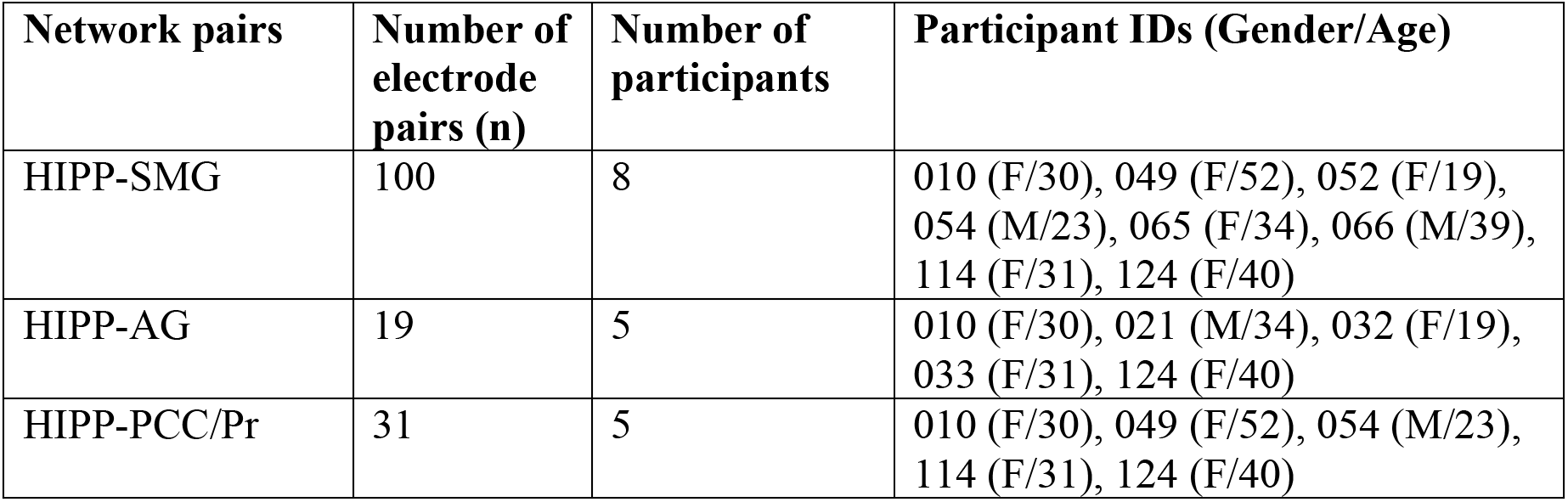
Number of electrode pairs used in phase transfer entropy (PTE) analysis in the spatial memory task. HIPP: hippocampus; SMG: supramarginal gyrus; AG: angular gyrus; PCC: posterior cingulate cortex, Pr: precuneus.

**Table 3c.** Number of electrodes in each node used in high-gamma power analysis in the spatial memory task. HIPP: hippocampus; SMG: supramarginal gyrus; AG: angular gyrus; PCC: posterior cingulate cortex, Pr: precuneus.

| Brain regions | Number of electrodes (n) | Number of participants | Participant IDs (Gender/Age) |
| --- | --- | --- | --- |
| HIPP | 31 | 12 | 010 (F/30), 021 (M/34), 032 (F/19), 033 (F/31), 049 (F/52), 052 (F/19), 054 (M/23), 065 (F/34), 066 (M/39), 089 (M/36), 114 (F/31), 124 (F/40) |
| SMG | 89 | 18 | 010 (F/30), 014 (F/47), 015 (F/54), 030 (M/23), 044 (M/58), 049 (F/52), 050 (M/20), 052 (F/19), 054 (M/23), 065 (F/34), 066 (M/39), 067 (F/45), 069 (M/26), 074 (M/24), 075 (M/50), 114 (F/31), 124 (F/40), 177 (F/23) |
| AG | 34 | 14 | 008 (F/55), 010 (F/30), 014 (F/47), 021 (M/34), 024 (F/36), 032 (F/19), 033 (F/31), 034 (F/29), 050 (M/20), 069 (M/26), 073 (M/60), 083 (F/49), 101 (F/26), 124 (F/40) |
| PCC/Pr | 42 | 13 | 006 (F/20), 010 (F/30), 018 (M/47), 024 (F/36), 030 (M/23), 044 (M/58), 049 (F/52), 051 (F/24), 054 (M/23), 074 (M/24), 101 (F/26), 114 (F/31), 124 (F/40) |

Participants were shown objects in various locations during the encoding periods and asked to retrieve the location of the objects during the recall period.

Surrogate data analysis revealed that causal influence from the hippocampus to the SMG, AG, and PCC/precuneus and the reverse were significantly higher than those expected by chance (*p*<0.05 in all cases) during the encoding periods in the SM task, indicating bidirectional causal influence between the hippocampus and the parietal cortex during memory encoding **(Figure 2c).** Similar to the verbal free recall tasks, the hippocampus had higher causal influences on SMG (*F*(1, 192) = 151.77, *p*<0.001, Cohen’s *d* = 1.78), AG (*F*(1, 32) = 17.63, *p*<0.001, Cohen’s *d* = 1.48), and PCC/precuneus (*F*(1, 56) = 199.58, *p*<0.001, Cohen’s *d* = 3.76) than the reverse, during spatial memory encoding (**Figure 3c**).

### Causal influences from the hippocampus to parietal cortex in delta-theta band during memory recall

#### VFR task

Surrogate data analysis revealed that causal influence from the hippocampus to the SMG, AG, and PCC/precuneus and the reverse were significantly higher than those expected by chance (*p*<0.05 in all cases) during the recall periods in the VFR task, indicating bidirectional causal influence between the hippocampus and the parietal cortex during memory recall **(Figure 2a).**

Moreover, the hippocampus had higher causal influences on SMG (*F*(1, 342) = 143.55, *p*<0.001, Cohen’s *d* = 1.30), AG (*F*(1, 130) = 70.70, *p*<0.001, Cohen’s *d* = 1.48), and PCC/precuneus (*F*(1, 93) = 45.80, *p*<0.001, Cohen’s *d* = 1.40) than the reverse, during memory recall (**Figure 3a**).

#### CATVFR task

Causal influence from the hippocampus to the SMG, AG, and PCC/precuneus and the reverse were significantly higher than those expected by chance (*p*<0.05 in all cases) during the recall periods in the CATVFR task, indicating bidirectional causal influence between the hippocampus and the parietal cortex during memory recall **(Figure 2b).**

The hippocampus also had higher causal influences on SMG (*F*(1, 160) = 51.29, *p*<0.001, Cohen’s *d* = 1.13), AG (*F*(1, 89) = 55.81, *p*<0.001, Cohen’s *d* = 1.58), and PCC/precuneus (*F*(1, 57) = 53.37, *p*<0.001, Cohen’s *d* = 1.93) than the reverse, during memory recall (**Figure 3b**).

#### SM task

Causal influence from the hippocampus to the SMG, AG, and PCC/precuneus and the reverse were significantly higher than those expected by chance (*p*<0.05 in all cases) during the recall periods in the SM task, indicating bidirectional causal influence between the hippocampus and the parietal cortex during spatial memory recall **(Figure 2c).**

Similar to the verbal free recall tasks, the hippocampus had higher causal influences on SMG (*F*(1, 192) = 157.25, *p*<0.001, Cohen’s *d* = 1.81), AG (*F*(1, 33) = 16.94, *p*<0.001, Cohen’s *d* = 1.44), and PCC/precuneus (*F*(1, 57) = 160.71, *p*<0.001, Cohen’s *d* = 3.36) than the reverse, during spatial memory recall (**Figure 3c**).

These results demonstrate a key role for delta-theta frequency signaling underlying higher causal influences of the hippocampus on the parietal cortex during both memory encoding and recall conditions.

### Replication of causal directed connectivity between the hippocampus and the parietal cortex across memory domains in the delta-theta band

We used replication BF analysis to estimate the degree of replicability of direction of causal influence across multiple tasks and domains (**Table 4**). Specifically, we used the posterior distribution obtained from the VFR (original) dataset as a prior distribution for the test of data from the CATVFR and SM (replication) datasets (Ly et al., 2019) (see **Methods** for details).

**Table 4.**
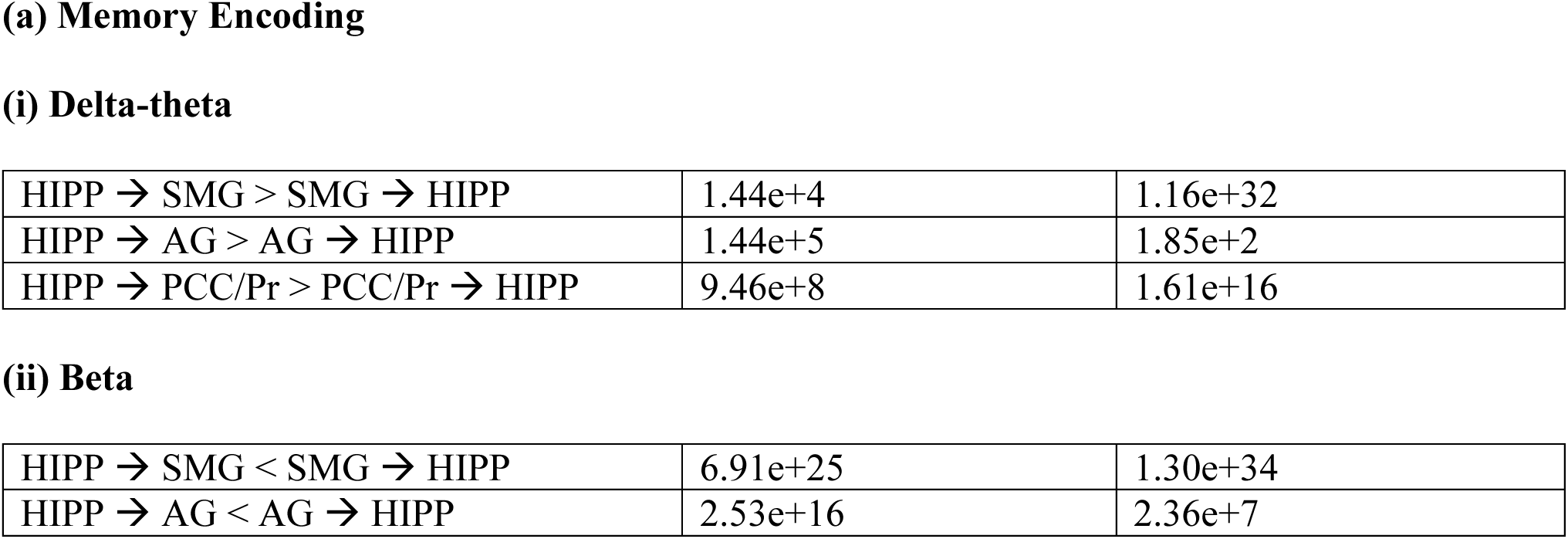
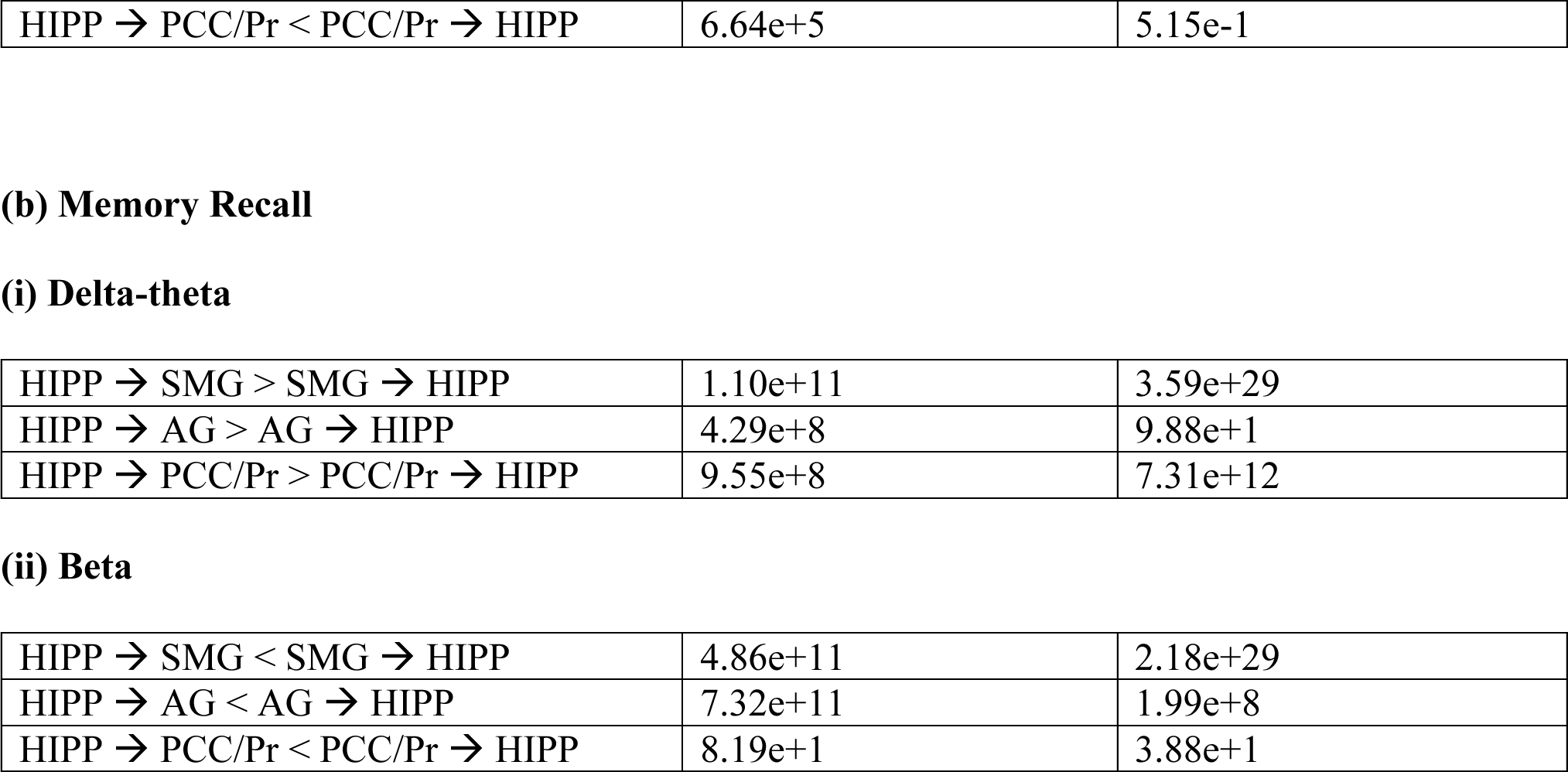
Replicability of findings of causal interactions of the hippocampus with SMG, AG, and PCC/Pr across verbal free recall, categorized verbal free recall, and spatial memory domains. (a) Memory Encoding and (b) Memory Recall. The verbal free recall (VFR) task was considered the original dataset and the categorized verbal free recall (CATVFR) and spatial memory (SM) tasks were considered replication datasets and Bayes factor (BF) for replication was calculated for pairwise tasks (verbal free recall vs. T, where T can be categorized verbal free recall, or spatial memory task). SMG: supramarginal gyrus, AG: angular gyrus, PCC: posterior cingulate cortex, Pr: precuneus.

(i) Delta-theta
|  |  |  |
| --- | --- | --- |
| HIPP → SMG > SMG → HIPP | 1.44e+4 | 1.16e+32 |
| HIPP → AG > AG → HIPP | 1.44e+5 | 1.85e+2 |
| HIPP → PCC/Pr > PCC/Pr → HIPP | 9.46e+8 | 1.61e+16 |
| (ii) Beta |  |  |
| HIPP → SMG < SMG → HIPP | 6.91e+25 | 1.30e+34 |
| HIPP → AG < AG → HIPP | 2.53e+16 | 2.36e+7 |
| HIPP → PCC/Pr < PCC/Pr → HIPP | 6.64e+5 | 5.15e-1 |
| (i) Delta-theta |  |  |
| HIPP → SMG > SMG → HIPP | 1.10e+11 | 3.59e+29 |
| HIPP → AG > AG → HIPP | 4.29e+8 | 9.88e+1 |
| HIPP → PCC/Pr > PCC/Pr → HIPP | 9.55e+8 | 7.31e+12 |
| (ii) Beta |  |  |
| HIPP → SMG < SMG → HIPP | 4.86e+11 | 2.18e+29 |
| HIPP → AG < AG → HIPP | 7.32e+11 | 1.99e+8 |
| HIPP → PCC/Pr < PCC/Pr → HIPP | 8.19e+1 | 3.88e+1 |

#### Encoding

Findings corresponding to the directed connectivity between the hippocampus and the SMG (BFs 1.44e+4 and 1.16e+32 for CATVFR and SM respectively), AG (BFs 1.44e+5 and 1.85e+2 for CATVFR and SM respectively), and PCC/precuneus (BFs 9.46e+8 and 1.61e+16 for CATVFR and SM respectively) during memory encoding were replicated in the delta-theta band across tasks (**Table 4a**).

#### Recall

Findings corresponding to the directed connectivity between the hippocampus and the SMG (BFs 1.10e+11 and 3.59e+29 for CATVFR and SM respectively), AG (BFs 4.29e+8 and 9.88e+1 for CATVFR and SM respectively), and PCC/precuneus (BFs 9.55e+8 and 7.31e+12 for CATVFR and SM respectively) during memory recall were also replicated in the delta-theta band across tasks (**Table 4b**).

These results demonstrate very high replicability of directed causal influences from the hippocampus to the parietal cortex nodes in the delta-theta frequency band, during both memory encoding and recall conditions.

### Causal influence from the hippocampus to the parietal cortex in the beta frequency band during memory encoding

Next, we examined directed connectivity between the hippocampus and parietal cortex in the beta frequency (12-30 Hz) band based on emerging findings in non-human primates regarding cortical signaling in the beta band during cognition (Engel & Fries, 2010) and based on our recent findings where we found higher directed causal influence from the prefrontal cortex to the hippocampus than the reverse, in beta frequency band during both encoding and recall, and across verbal and spatial episodic memory domains. We computed PTE from the parietal cortex nodes to the hippocampus, and in the reverse direction, during encoding and recall in the beta frequency band.

#### VFR task

Surrogate data analysis revealed that causal influence from the hippocampus to the SMG, AG, and PCC/precuneus and the reverse were significantly reduced compared to those expected by chance (*p*<0.05 in all cases) (**Figure 4a**). Crucially, SMG (*F*(1, 343) = 249.25, *p*<0.001, Cohen’s *d* = 1.71), AG (*F*(1, 129) = 167.3, *p*<0.001, Cohen’s *d* = 2.28), and PCC/precuneus (*F*(1, 92) = 40.78, *p*<0.001, Cohen’s *d* = 1.33) had higher causal influences on the hippocampus than the reverse, during memory encoding (**Figure 5a**).

**Figure 4.**
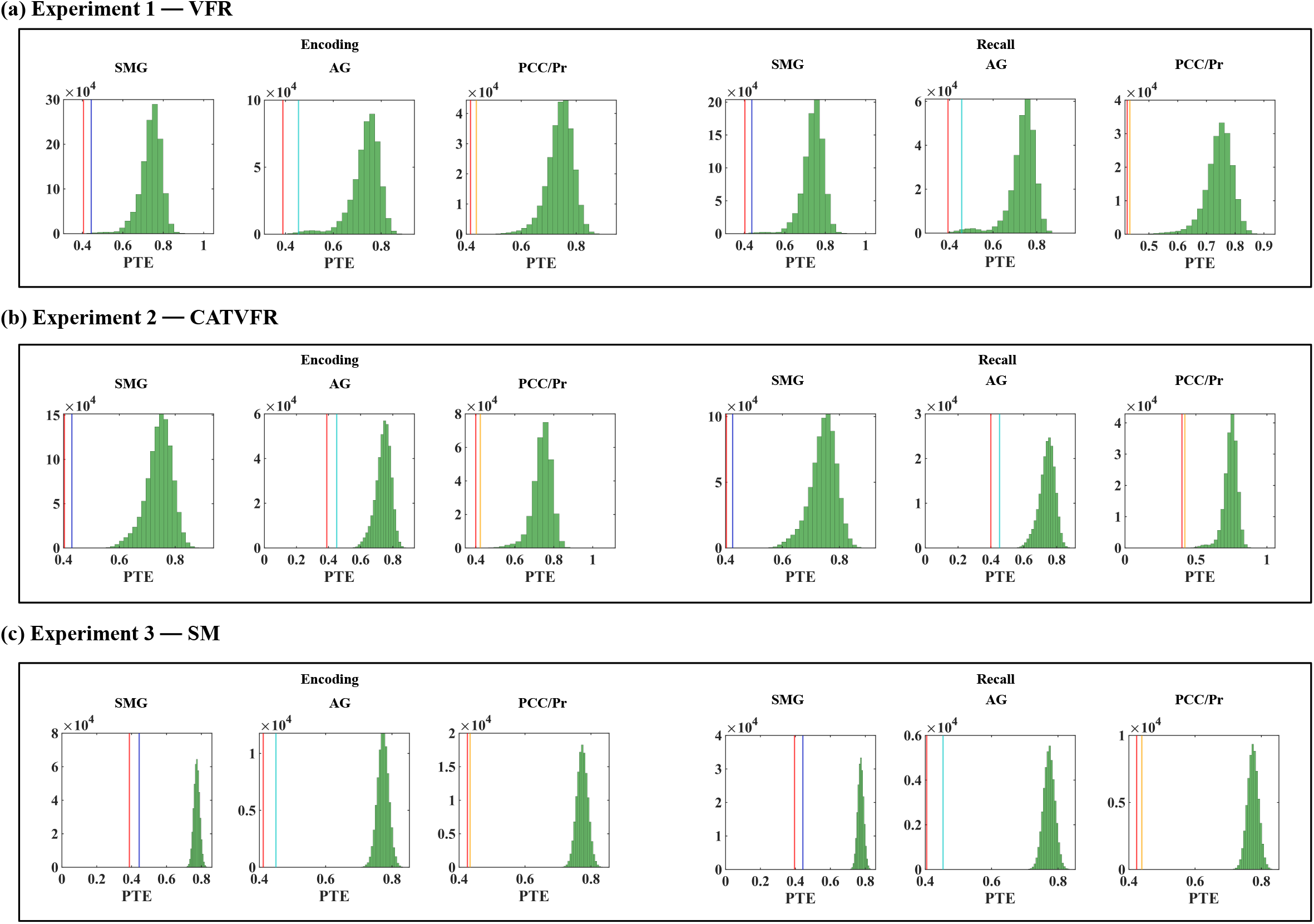
Surrogate data analysis results related to directed causal influence from the hippocampus to the parietal cortex (red) and the reverse (blue for supramarginal gyrus, cyan for angular gyrus, and orange for PCC/precuneus) in the beta band, during (a) VFR, (b) CATVFR, and (c) SM experiments. Surrogate data analysis revealed that causal influence from the hippocampus to the SMG, AG, and PCC/precuneus and the reverse were significantly reduced compared to those expected by chance during both encoding and recall periods, and across the three experiments (all *ps* < 0.05).

**Figure 5.**
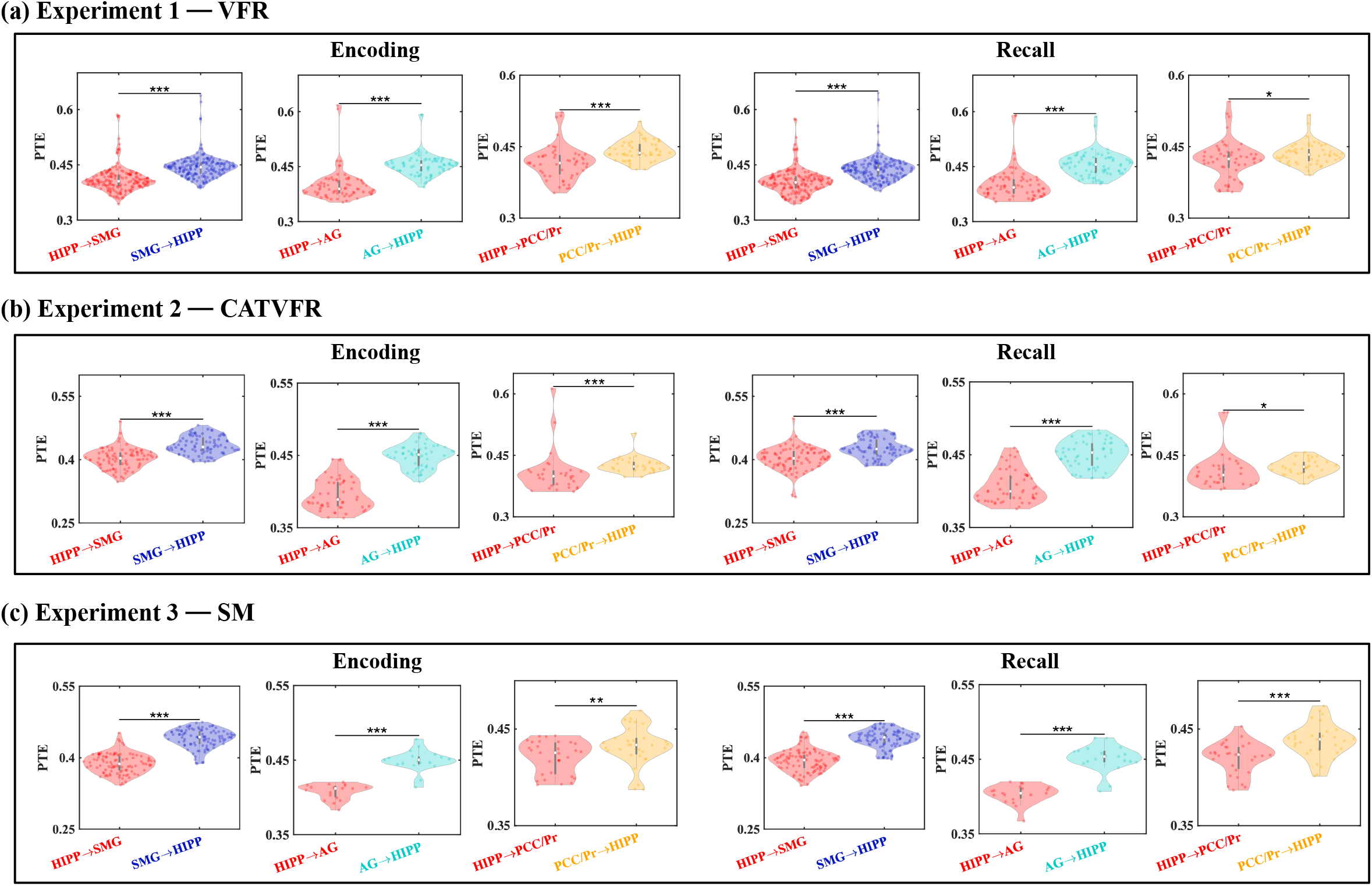
Directed connectivity between hippocampus and SMG, AG, and PCC/precuneus in beta band (12-30 Hz). **(a) Experiment 1, VFR:** SMG (SMG → HIPP) (n=184), AG (AG → HIPP) (n=72), and PCC/precuneus (PCC/Pr → HIPP) (n=53) showed higher causal influences on the hippocampus during both memory encoding and recall, compared to the reverse direction (HIPP → SMG, HIPP → AG, and HIPP → PCC/Pr respectively). **(b) Experiment 2, CATVFR:** SMG (SMG → HIPP) (n=87), AG (AG → HIPP) (n=50), and PCC/precuneus (PCC/Pr → HIPP) (n=31) showed higher causal influences on the hippocampus during both memory encoding and recall, compared to the reverse direction (HIPP → SMG, HIPP → AG, and HIPP → PCC/Pr respectively). **(c) Experiment 3, SM:** SMG (SMG → HIPP) (n=100), AG (AG → HIPP) (n=19), and PCC/precuneus (PCC/Pr → HIPP) (n=31) also showed higher causal influences on the hippocampus during both memory encoding and recall, compared to the reverse direction (HIPP → SMG, HIPP → AG, and HIPP → PCC/Pr respectively). *** *p* < 0.001, ** *p* < 0.01, * *p* < 0.05 (Linear mixed effects ANOVA, FDR-corrected).

#### CATVFR task

Surrogate data analysis revealed that causal influence from the hippocampus to the SMG, AG, and PCC/precuneus and the reverse were significantly reduced compared to those expected by chance (*p*<0.05 in all cases) (**Figure 4b**). Crucially, SMG (*F*(1, 161) = 132.77, *p*<0.001, Cohen’s *d* = 1.82), AG (*F*(1, 98) = 142.37, *p*<0.001, Cohen’s *d* = 2.41), and PCC/precuneus (*F*(1, 57) = 29.49, *p*<0.001, Cohen’s *d* = 1.44) had higher causal influences on the hippocampus than the reverse during memory encoding (**Figure 5b**).

#### SM task

Surrogate data analysis revealed that causal influence from the hippocampus to the SMG, AG, and PCC/precuneus and the reverse were significantly reduced compared to those expected by chance in the SM task (*p*<0.05 in all cases) (**Figure 4c**). Crucially, similar to the verbal free recall tasks, SMG (*F*(1, 191) = 276.92, *p*<0.001, Cohen’s *d* = 2.41), AG (*F*(1, 32) = 73.29, *p*<0.001, Cohen’s *d* = 3.01), and PCC/precuneus (*F*(1, 56) = 7.39, *p*<0.01, Cohen’s *d* = 0.73) had higher causal influences on the hippocampus than the reverse, during spatial memory encoding (**Figure 5c**).

### Causal influence from the hippocampus to the parietal cortex in the beta frequency band during memory recall

#### VFR task

Surrogate data analysis revealed that causal influence from the hippocampus to the SMG, AG, and PCC/precuneus and the reverse were significantly reduced compared to those expected by chance (*p*<0.05 in all cases) (**Figure 4a**). Crucially, SMG (*F*(1, 344) = 225.68, *p*<0.001, Cohen’s *d* = 1.62), AG (*F*(1, 129) = 186.1, *p*<0.001, Cohen’s *d* = 2.40), and PCC/precuneus (*F*(1, 92) = 6.25, *p*<0.05, Cohen’s *d* = 0.52) had higher causal influences on the hippocampus than the reverse, during memory recall (**Figure 5a**).

#### CATVFR task

Causal influence from the hippocampus to the SMG, AG, and PCC/precuneus and the reverse were significantly reduced compared to those expected by chance (*p*<0.05 in all cases) (**Figure 4b**). Crucially, SMG (*F*(1, 160) = 52.84, *p*<0.001, Cohen’s *d* = 1.15), AG (*F*(1, 98) = 114.09, *p*<0.001, Cohen’s *d* = 2.16), and PCC/precuneus (*F*(1, 57) = 6.51, *p*<0.05, Cohen’s *d* = 0.67) had higher causal influences on the hippocampus than the reverse, during memory recall (**Figure 5b**).

#### SM task

Causal influence from the hippocampus to the SMG, AG, and PCC/precuneus and the reverse were significantly reduced compared to those expected by chance (*p*<0.05 in all cases) (**Figure 4c**). Crucially, similar to the verbal free recall tasks, SMG (*F*(1, 192) = 230.41, *p*<0.001, Cohen’s *d* = 2.19), AG (*F*(1, 32) = 61.55, *p*<0.001, Cohen’s *d* = 2.79), and PCC/precuneus (*F*(1, 54) = 20.03, *p*<0.001, Cohen’s *d* = 1.22) had higher causal influences on the hippocampus than the reverse, during spatial memory recall (**Figure 5c**).

These results demonstrate that causal influences of the parietal cortex on the hippocampus are relatively enhanced, compared to the reverse, during both memory encoding and recall conditions in each of the three tasks.

### Replication of causal directed connectivity between the hippocampus and the parietal cortex across memory domains in the beta band

We next repeated the replication BF analysis to estimate the degree of replicability of directed connectivity across multiple tasks and memory domains in the beta frequency band (**Table 4**).

#### Encoding

Findings corresponding to the directed connectivity between the hippocampus and the SMG (BFs 6.91e+25 and 1.30e+34 for CATVFR and SM respectively) and AG (BFs 2.53e+16 and 2.36e+7 for CATVFR and SM respectively) were replicated in the beta band across tasks (**Table 4a**). Findings corresponding to the directed connectivity between the hippocampus and the PCC/precuneus was replicated for the CATVFR task in the beta band (BF 6.64e+5) (**Table 4a**).

#### Recall

Findings corresponding to the directed connectivity between the hippocampus and the SMG (BFs 4.86e+11 and 2.18e+29 for CATVFR and SM respectively), AG (BFs 7.32e+11 and 1.99e+8 for CATVFR and SM respectively), and PCC/precuneus (BFs 8.19e+1 and 3.88e+1 for CATVFR and SM respectively) during memory recall were also replicated in the beta band across tasks (**Table 4b**).

Together, these results demonstrate very high replicability of relatively enhanced directed connectivity from the lateral and medial parietal cortex nodes to the hippocampus in the beta frequency band, during both memory encoding and recall conditions.

### Directed causal influence from the SMG to the hippocampus during encoding influences memory performance

We examined whether the directed connectivity between the hippocampus and the parietal cortex is dependent on memory performance. For this, we estimated the differential causal influence for the high and low memory performers. We defined high and low memory performers based on whether their recall accuracy was higher or lower than the median recall accuracy, respectively.

#### VFR task

We found that causal influence from the SMG to the hippocampus increased for the high compared to low memory performers during memory encoding (*F*(1, 170) = 28.60, *p*<0.001, Cohen’s *d* = 0.82) in the delta-theta frequency band (**Figure 6a**).

**Figure 6.**
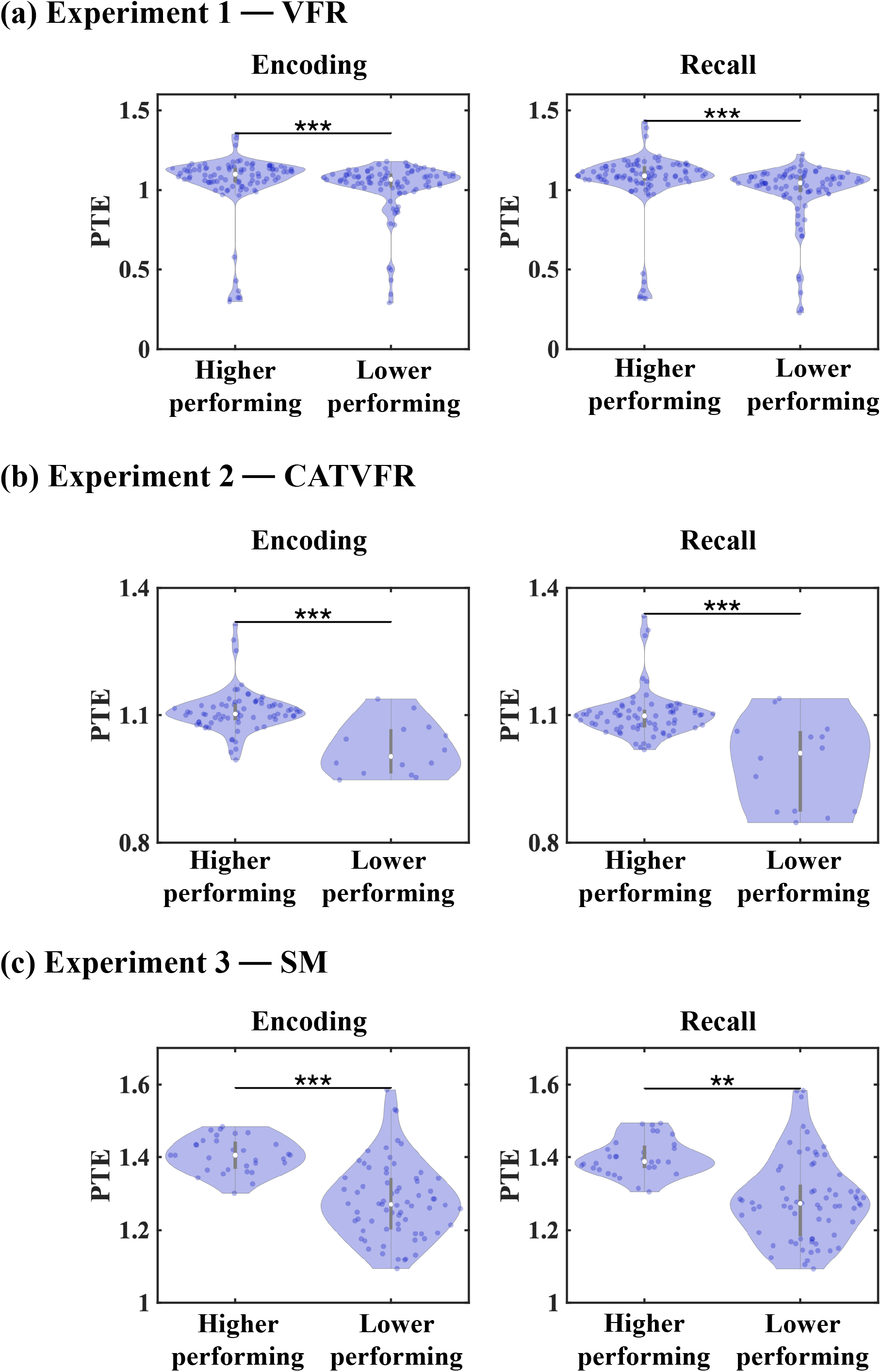
SMG → HIPP directed causal influence increased for the higher compared to lower memory performers in the delta-theta band, during (a) VFR, (b) CATVFR, and (c) SM experiments. *** *p* < 0.001, ** *p* < 0.01 (Linear mixed effects ANOVA, FDR-corrected).

#### CATVFR task

Causal influence from the SMG to the hippocampus also increased for the high compared to low memory performers during memory encoding in the CATVFR task (*F*(1, 81) = 32.19, *p*<0.001, Cohen’s *d* = 1.26) in the delta-theta band (**Figure 6b**).

#### SM task

Causal influence from the SMG to the hippocampus increased for the high compared to low memory performers during spatial memory encoding (*F*(1, 75) = 21.05, *p*<0.001, Cohen’s *d* = 1.06) in the delta-theta band (**Figure 6c**).

We also used replication BFs to assess the degree of replicability of the differential causal influence from the SMG to the hippocampus for the high compared to the low performers. This analysis revealed that the differential causal influence for the high compared to low performers was replicated for the encoding periods of the memory tasks (BFs 9.78e+6 and 8.39e+8 for CATVFR and SM respectively).

We did not find any consistent differential PTE signature for the high compared to the low memory performers in any other frequency band or brain region, highlighting a critical role of the SMG causal influence on the hippocampus for memory performance in the delta-theta frequency band during memory encoding.

### Directed causal influence from the SMG to the hippocampus during recall influences memory performance

#### VFR task

We found that causal influence from the SMG to the hippocampus increased for the high compared to low memory performers during memory recall (*F*(1, 170) = 44.07, *p*<0.001, Cohen’s *d* = 1.02) in the VFR task in the delta-theta frequency band (**Figure 6a**).

#### CATVFR task

Causal influence from the SMG to the hippocampus increased for the high compared to low memory performers in the CATVFR task (*F*(1, 81) = 33.99, *p*<0.001, Cohen’s *d* = 1.30) in the delta-theta frequency band (**Figure 6b**).

#### SM task

Causal influence from the SMG to the hippocampus increased for the high compared to low memory performers during spatial memory recall (*F*(1, 88) = 9.15, *p*<0.01, Cohen’s *d* = 0.64) in the delta-theta band (**Figure 6c**).

We also used replication BFs to assess the degree of replicability of the differential causal influence from the SMG to the hippocampus for the high compared to the low performers during the recall periods. This analysis revealed that the differential causal influence for the high compared to low performers was replicated for the recall periods of the memory tasks (BFs 1.47e+6 and 5.45e+8 for CATVFR and SM respectively). Together, these results demonstrate robust replicability of higher directed causal influence from the SMG to the hippocampus for the high memory performers compared to the low performers in the delta-theta band, during both memory encoding and recall.

We did not find any consistent differential PTE signature for the high compared to the low memory performers in any other frequency band or brain region during the recall periods, highlighting a critical role of the SMG causal influence on the hippocampus for memory performance in the delta-theta frequency band.

### Differential causal directed connectivity between the hippocampus and the parietal cortex for the correctly compared to incorrectly recalled trials

We further examined memory effects by comparing PTE between correct and incorrect trials. However, this analysis did not reveal differences in causal influence from the hippocampus to the SMG, AG, and PCC/precuneus or the reverse, between correctly and incorrectly recalled trials during the encoding as well as recall periods of the verbal or spatial memory tasks in any frequency band (all *ps*>0.05).

### Task-related high-gamma power in hippocampus compared to SMG, AG, and PCC/precuneus during encoding and recall

We next examined neuronal activity in the hippocampus and the SMG, AG, and PCC/precuneus and tested whether activity in the PCC/precuneus is suppressed compared to activity in the hippocampus. Previous studies have suggested that power in the high gamma band (80-160 Hz) is correlated with fMRI BOLD signals (Hutchison, Hashemi, Gati, Menon, & Everling, 2015; Lakatos, Gross, & Thut, 2019; Leopold, Murayama, & Logothetis, 2003; Mantini, Perrucci, Del Gratta, Romani, & Corbetta, 2007; Scholvinck, Maier, Ye, Duyn, & Leopold, 2010), and is thought to reflect local neuronal activity (Canolty & Knight, 2010). Therefore, we compared high-gamma power (**Methods**) in the hippocampus with the SMG, AG, and PCC/precuneus during memory encoding and recall conditions across verbal and spatial episodic memory domains.

This analysis revealed that high-gamma power in the PCC/precuneus was suppressed compared to high-gamma power in the hippocampus almost throughout the encoding periods of the verbal free recall, categorized verbal free recall, and spatial memory tasks (*ps*<0.05) (**Figures 7a-c**). High-gamma power in the SMG was also suppressed compared to high-gamma power in the hippocampus during the encoding periods of the verbal free recall, categorized verbal free recall, and spatial memory tasks (*ps*<0.05) (**Figures 7a-c**), although this effect was less consistent compared to PCC/precuneus. High-gamma power results during the recall periods were not consistent across the three tasks (**Figures 7a-c**). Moreover, high-gamma power of the hippocampus compared to those from the AG was also not consistent across the three tasks (**Figures 7a-c**).

**Figure 7.**
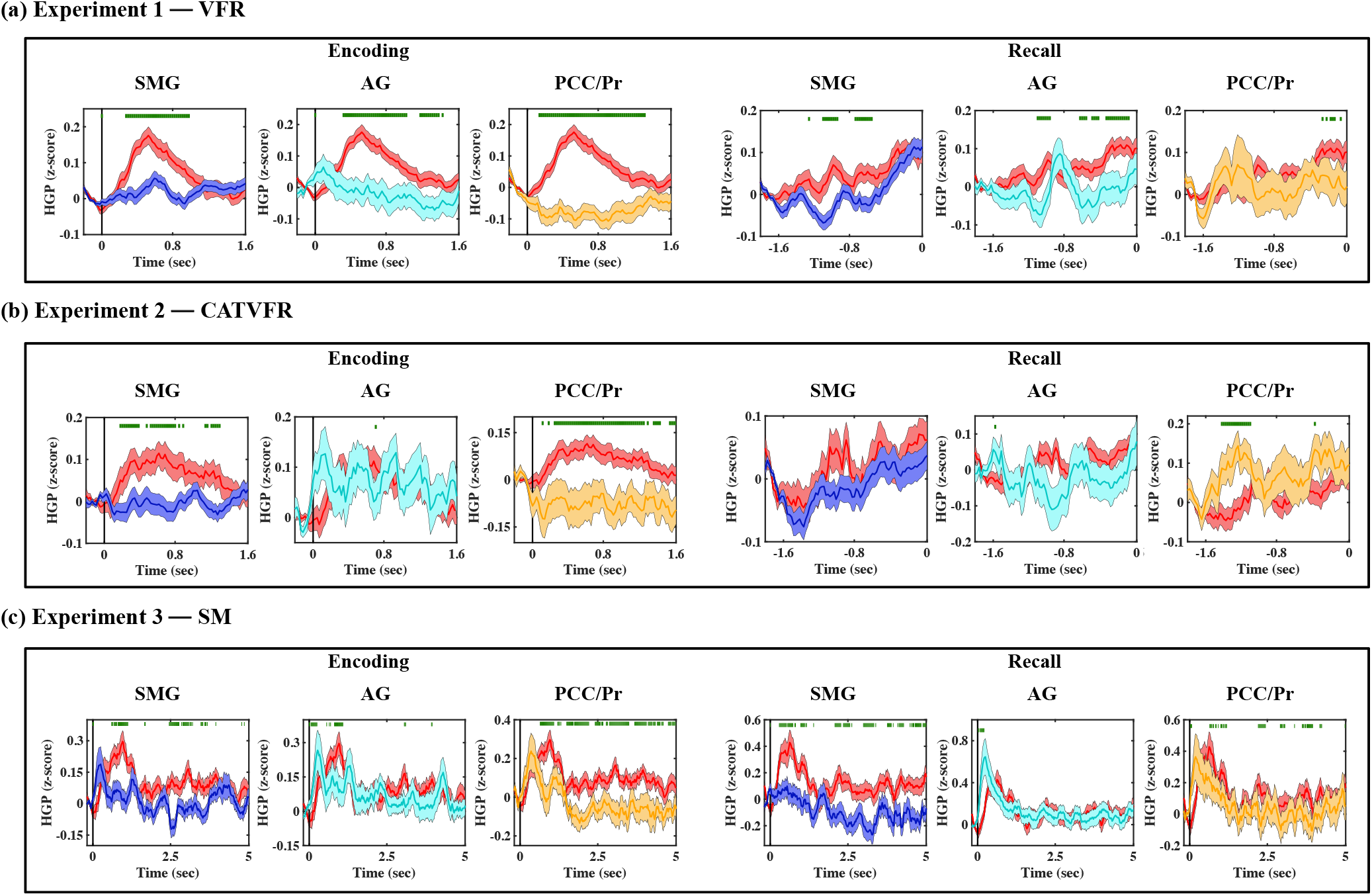
iEEG evoked response, quantified using high-gamma power (HGP), for HIPP (red) and SMG (blue), AG (cyan), and PCC/precuneus (orange) during (a) VFR, (b) CATVFR, and (c) SM experiments. Horizontal green lines denote *ps* < 0.05.

## Discussion

Our investigation focused on the electrophysiological underpinnings of directed connectivity between the hippocampus and the parietal cortex during three different cognitive tasks involving verbal and spatial memory processing in humans. Utilizing iEEG recordings from a substantial cohort of 96 individuals, we dissected the frequency-dependent nature of interactions between the hippocampus and parietal cortex. Our analysis revealed a consistent pattern of higher causal directed connectivity in the delta-theta band from the hippocampus to both the SMG and AG subdivisions of the lateral parietal cortex as well as the PCC/precuneus subdivisions of the medial parietal cortex. This directional connectivity was observed during both memory encoding and recall in each of the three tasks, with high replication Bayes Factor (BF). Crucially, directed connectivity from the SMG to the hippocampus was enhanced in participants with higher, compared to lower memory recall, highlighting the behavioral relevance of delta-theta band-specific directed connectivity between the hippocampus and the parietal cortex. Our findings provide novel insights into asymmetric frequency-specific directed connectivity between the hippocampus and the parietal cortex during verbal and spatial information processing in the human brain (**Figure 8**).

**Figure 8.**
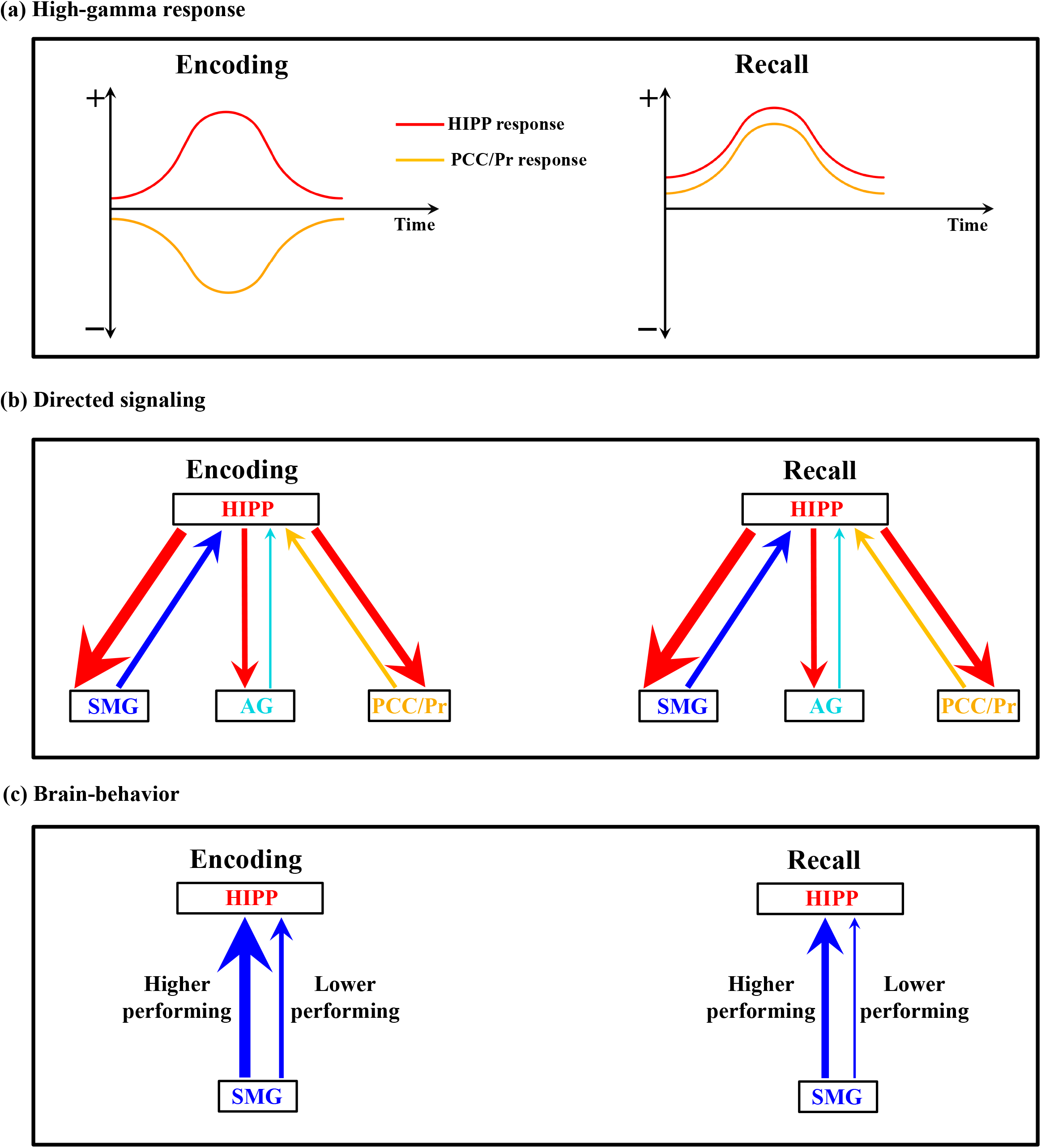
Schematic illustration of key findings related to the intracranial electrophysiology of directed connectivity between the hippocampus and the lateral and medial parietal cortex in human episodic memory. **(a) High-gamma response:** Our analysis of local neuronal activity revealed consistent suppression of high-gamma power in the PCC/precuneus compared to the hippocampus during encoding periods across all three episodic memory experiments. In contrast, we detected similar high-gamma band power in the PCC/precuneus relative to the hippocampus during the recall periods. **(b) Directed signaling:** We found stronger causal influence (denoted by red arrows, thickness of arrows denotes degree of replicability across experiments, see **Table 4**) by the hippocampus on the lateral and medial parietal cortex than the reverse, in the delta-theta band, across all three memory experiments, and during both encoding and recall periods. **(c) Brain-behavior:** Brain-behavior analysis revealed that SMG → HIPP directed causal influence increased for the higher compared to lower memory performers in the delta-theta band, across all three memory experiments, and during both encoding and recall periods.

### Frequency specific directed connectivity between hippocampus and parietal cortex

Our analysis of causal signaling was based on phase transfer entropy (PTE), which provides a robust and powerful tool for characterizing directed connectivity between brain regions based on phase coupling (Hillebrand et al., 2016; Lobier et al., 2014; M. Y. Wang et al., 2017). Previous findings using multielectrode array recordings in both humans and animal models have established that phase, rather than amplitude, is crucial for both spatial and temporal encoding of information in the brain (Kayser, Montemurro, Logothetis, & Panzeri, 2009; Lachaux, Rodriguez, Martinerie, & Varela, 1999; Lopour, Tavassoli, Fried, & Ringach, 2013; Ng, Logothetis, & Kayser, 2013; Siegel, Warden, & Miller, 2009). PTE assesses with the ability of one time-series to predict future values of other time-series thus estimating the time-delayed causal influences between the two time-series. Crucially, PTE is a robust, nonlinear measure of directed connectivity between time-series (Das, de Los Angeles, & Menon, 2022; Das & Menon, 2020, 2021, 2022a, 2022b, 2023; Hillebrand et al., 2016; Lobier et al., 2014).

Electrophysiological studies in rodents have reported strong delta (0.5-4 Hz), theta (4-8 Hz), and beta (12-30 Hz) frequency band oscillations in the hippocampus (Schultheiss et al., 2020; Trimper, Galloway, Jones, Mandi, & Manns, 2017; Trimper, Stefanescu, & Manns, 2014) and theta-gamma phase-amplitude coupling between the hippocampus and the parietal cortex, with hippocampus-theta modulating gamma power in the parietal cortex (Sirota et al., 2008). Based on previous reports in rodents, non-human primates, and humans, we focused on delta-theta (0.5-8 Hz) and beta (12-30 Hz), frequency bands, as enhanced local field potentials in these frequency bands have been identified in the hippocampus and parietal cortex (Ekstrom et al., 2005; Ekstrom & Watrous, 2014; Engel & Fries, 2010; Gonzalez et al., 2015; Watrous et al., 2013). In our previous iEEG studies, we have found higher causal influence from the hippocampus to the prefrontal cortex than the reverse, in delta-theta frequency band and higher causal influence from the prefrontal cortex to the hippocampus than the reverse, in beta frequency band, during both encoding and recall, and across verbal and spatial episodic memory domains (Das & Menon, 2021, 2022b). Moreover, previous iEEG studies have reported significant delta-theta frequency band activity in the hippocampus during recall of spatial information from recently encoded memories (Goyal et al., 2018; Jacobs et al., 2016), but the common frequency-specificity of causal hippocampal-parietal cortex signaling in the human brain associated with verbal and spatial memory encoding and recall has not been well understood.

PTE revealed significantly higher causal influence of the hippocampal electrodes on the parietal cortex electrodes than the reverse, during both encoding and recall of memory in the delta-theta band. Crucially, this asymmetric pattern of directed connectivity was replicated across both verbal and spatial memory tasks. These findings build upon our recent iEEG findings where we have shown replicable pattern of directed causal influence from the hippocampus to the middle frontal gyrus and inferior frontal gyrus subdivisions of the prefrontal cortex than the reverse, during memory encoding and recall and across verbal and spatial memory domains (Das & Menon, 2021, 2022b). The current findings thus suggest that the hippocampus has directed causal influence on broad cortical areas involving both the prefrontal and parietal cortices. Our findings also converge on recent findings from iEEG recordings in humans that have shown that the left hemisphere hippocampus plays a crucial role for episodic and spatial memory processing (Ekstrom et al., 2005; Gonzalez et al., 2015). Furthermore, recent iEEG studies involving intracranial stimulation of the bilateral medial temporal lobe suggest that directed connectivity between hippocampus and parietal cortex contributes to successful memory retrieval (Das & Menon, 2023). Results are also consistent with previous multiple structural and functional MRI (Grön, Wunderlich, Spitzer, Tomczak, & Riepe, 2000; Iglói, Doeller, Berthoz, Rondi-Reig, & Burgess, 2010; Kaplan, Horner, Bandettini, Doeller, & Burgess, 2014; Maguire, Burgess, & O’Keefe, 1999; Maguire et al., 2000), positron imaging tomography (PET) (Ghaem et al., 1997; Maguire et al., 1998; Maguire et al., 1999), and magnetoencephalography (MEG) (Pu, Cornwell, Cheyne, & Johnson, 2017) studies in humans which have revealed the involvement of bilateral hippocampus in spatial navigation and memory. Our findings thus provide robust electrophysiological evidence for dynamic causal influence of the left hippocampus on both the lateral and medial parietal cortex during both verbal and spatial episodic memory tasks.

### Left hemisphere hippocampus plays a critical role in both verbal and spatial memory processing

We focused on dynamic interactions between the left hemisphere hippocampus and the parietal cortex. This focus was necessitated by the limitation that our iEEG data had inadequate coverage in the right hippocampus. While the left hemisphere hippocampus is widely recognized for its dominant role in verbal episodic memory encoding and recall (Burgess et al., 2002), its role in spatial memory processing has remained poorly understood. The contribution of the left hemisphere to spatial memory is less clear, although some studies point to the involvement of both the left and right hippocampi (Bohbot et al., 1998; Maguire et al., 1996; Spiers et al., 2001). These findings are supported by structural and functional MRI, PET, MEG, and iEEG studies, which have also implicated both the left and right hippocampi in spatial navigation and memory (Bohbot et al., 1998; Maguire, Burke, Phillips, & Staunton, 1996; Spiers et al., 2001). iEEG studies in humans have also pointed to the involvement of both the left and right hemisphere hippocampus in spatial navigation and memory (J. Miller et al., 2018; Stangl et al., 2021).

Importantly, our findings of left hippocampal causal signaling during both verbal and spatial memory aligns with recent work by Seger and colleagues, which suggests that memory might be a more potent driver of hippocampal theta oscillations in humans than navigation (Seger, Kriegel, Lega, & Ekstrom, 2023). In their study, patients who mentally simulated routes they had just navigated showed greater hippocampal theta power, higher frequency, and longer duration of oscillations than during actual navigation. This suggests that in humans, unlike in rodents, memory might be a more potent catalyst for hippocampal theta oscillations. These findings support models of internally generated theta oscillations in the human hippocampus and could indicate that memory, rather than spatial navigation, is the primary function driving hippocampal activity (Ekstrom & Hill, 2023). In our study, the focus on the left hemisphere, which is traditionally predominantly associated with verbal memory, assumes greater relevance in light of these new findings. It raises the possibility that the left hippocampus may play a more comprehensive role during memory processing, extending beyond verbal tasks to encompass spatial memory tasks as well.

### Directed connectivity and high-gamma power

The PCC/precuneus and the angular gyrus, two key nodes of the DMN, are typically deactivated during attention demanding tasks (Wen, Liu, Yao, & Ding, 2013). Despite this, they are directly implicated in episodic memory functions (Buckner et al., 2008; Menon, 2023). Human iEEG studies have shown a prominent role of the PCC/precuneus in episodic memory retrieval (Aponik-Gremillion et al., 2022; Foster, Dastjerdi, & Parvizi, 2012; Foster, Kaveh, Dastjerdi, Miller, & Parvizi, 2013; Foster, Rangarajan, Shirer, & Parvizi, 2015; Lega, Germi, & Rugg, 2017; Natu et al., 2019). In contrast, the supramarginal gyrus, a central node of the frontoparietal control network and integral to cognitive control over memory, typically displays heightened activity during memory tasks (Badre, Poldrack, Paré-Blagoev, Insler, & Wagner, 2005; Badre & Wagner, 2007; Wagner, Pare-Blagoev, Clark, & Poldrack, 2001; Wagner et al., 2005). Our high-resolution iEEG analysis revealed that, specifically during encoding, high-gamma power was consistently suppressed in the PCC/precuneus relative to the hippocampus across all three episodic memory experiments. This suppression was unique to the PCC/precuneus; no such consistent suppression pattern emerged in either the supramarginal or angular gyrus. Our findings significantly extend this knowledge base to specific periods of memory formation, leveraging high-resolution iEEG recordings for temporal precision. Importantly, even though the hippocampus and the PCC/precuneus are both associated with the default mode network, we observed a clear dissociation of high-gamma dynamics between these two brain areas. The PCC/precuneus high-gamma activity was suppressed compared to the hippocampus during encoding, and both brain regions were similarly active during the recall periods of the tasks, suggesting context-specific dissociation between these DMN nodes.

Contrary to our initial predictions regarding the AG’s alignment with the default mode network, we found that neural responses in this region were not suppressed compared to the hippocampus during either encoding or recall. Instead, the AG displayed heightened neural activity. One explanation may lie in the functional heterogeneity of the AG, which has distinct cytoarchitectonic subregions – PGa and PGp – with divergent functional roles (Bellana, Ladyka-Wojcik, Lahan, Moscovitch, & Grady, 2023; Niu & Palomero-Gallagher, 2023; Rockland & Graves, 2023; Uddin et al., 2010; Wu et al., 2009). The former appears aligned with the SMG and the frontoparietal network, while the latter is more closely associated with the default mode network (Seghier, 2013, 2023; Seghier, Fagan, & Price, 2010). Future research with adequate subregion sampling is necessary to resolve questions around this functional heterogeneity and its alignment with the default mode network.

High-gamma (80-160 Hz) activity is a reliable indicator of localized neuronal activity, with observed elevations during a variety of cognitive tasks including working memory and episodic memory (Canolty et al., 2006; Crone et al., 2001; Daitch & Parvizi, 2018; Edwards et al., 2005; Lachaux et al., 2005; Mainy et al., 2007; Sederberg, Schulze-Bonhage, Madsen, Bromfield, Litt, et al., 2007; Tallon-Baudry et al., 2005). The observed suppression in high-gamma power in the PCC/precuneus compared to the hippocampus specifically during memory encoding, but not during recall, offers strong evidence for the selective functional downregulation of this region when processing external stimuli during the encoding period of episodic memory formation. Interestingly, despite these patterns of local modulation in high-gamma power, our analysis found a substantial causal influence exerted by the hippocampus on both the lateral and medial parietal cortex in the low-frequency delta-theta (0.5-8 Hz) band. However, this causal influence was notably absent in the high-gamma frequency range. This distinction aligns with current models positing that high-gamma activity is predominantly indicative of localized processing, whereas lower-frequency oscillations are more likely to mediate long-range network interactions (Bastos et al., 2015; Das et al., 2022; Das & Menon, 2020, 2021, 2022b, 2023; K. J. Miller et al., 2007). Our findings thus underscore the necessity of differentiating between high-gamma and low-frequency fluctuations, as these separate spectral domains reflect distinct physiological phenomena with specific roles in neural network function during episodic memory tasks.

In contrast, we found an opposite pattern of directed connectivity in the beta band with higher parietal cortex causal influences on the hippocampus, compared to the reverse direction; again, this pattern was observed during both memory encoding and recall periods and across verbal and spatial memory domains. This robust pattern held despite relative decreases in phase transfer entropy in the beta band during memory task performance when compared to the background surrogate distribution. This aligns with supression of beta band phase coupling, compared to background surrogate distribution in both iEEG and EEG studies (Gaillard et al., 2009; Hillebrand et al., 2016). Crucially, our findings are aligned with previous iEEG studies reporting higher causal influence from the hippocampus to the prefrontal cortex than the reverse, in delta-theta frequency band and higher causal influence from the prefrontal cortex to the hippocampus than the reverse, in beta frequency band, during both encoding and recall, and across verbal and spatial episodic memory domains (Das & Menon, 2021, 2022b). Top-down causal influence from the parietal cortex in the beta-band may contribute to attention-related transitions of latent neuronal ensembles into “active” memory representations (Spitzer & Haegens, 2017).

### Behavioral relevance of hippocampal-parietal cortex directed connectivity

Our results also offer new insights into the mechanisms of memory formation specifically. Examination of differential causal influence between lower-performing and higher-performing participants revealed that causal delta-theta frequency band influences from the SMG to the hippocampus increased in the higher-performing, compared to the lower-performing, participants. This finding held true during both the memory encoding and recall periods and across the verbal free recall, categorized verbal free recall, and spatial memory experiments. High replication BFs further demonstrated the robust replicability of higher directed causal influence from the SMG to the hippocampus for the higher-performing, compared to lower-performing, participants. We did not find any consistent differential PTE signature between the two groups of participants in any other frequency band or in directed interactions involving the AG or the PCC/precuneus. These results provide new insights into our understanding of functional circuitry of the SMG in episodic memory.

Our results extend prior findings related to differential neural responses in the SMG and AG by examining directional signaling between these regions and the hippocampus. Using iEEG, Rubinstein et al. previously demonstrated that spectral power changes related to successful memory encoding and retrieval (“subsequent memory effects”) were stronger in the SMG compared to the AG (Rubinstein et al., 2021). Moreover, recent transcranial magnetic stimulation studies and fMRI findings suggest an important role for the SMG in episodic memory encoding (Ciaramelli et al., 2020; Guidali, Pisoni, Bolognini, & Papagno, 2019). Consistent with the localization of memory-related spectral power changes to SMG in the Rubinstein study, increased functional coupling specifically originating from this region provides further evidence that it plays a specialized role in coordinating episodic memory processes. This aligns with the attention-to-memory model which proposes that attention plays a critical role in the encoding and retrieval of memories (Cabeza, 2008; Cabeza et al., 2008; Cabeza et al., 2011). According to this model, attention focuses cognitive resources on relevant information, enhancing the encoding of information into memory and facilitating its subsequent retrieval. Our findings highlight a specific role for the SMG in this process.

In contrast to the SMG, the AG did not show enhanced causal interactions in higher-relative to lower-performing participants. The AG is thought to facilitate recall when more complex binding of semantics or concepts is needed, processes that are less relevant to the memory tasks used in the present study. By demonstrating that the SMG ramps up directional signaling to hippocampal memory structures during encoding and retrieval among higher performing participants, the current results suggest that localized spectral power changes reported by Rubinstein and colleagues may reflect efficient engagement of memory control processes. Future work should explore how interactions between attention-related SMG signaling and conceptual binding mediated by AG underpin memory encoding and recall. Elucidating both representational and control processes across parietal subnetworks will shed further light on neurocomputational mechanisms of episodic memory.

### Reinterpreting findings in the context of attention and general cognitive processes

Our analysis revealed no significant differences in directed connectivity between correct and incorrect memory trials, suggesting that the reported effects may not be specific to successful memory formation. Instead, the differences found between high and low performers point to the possibility that these connectivity patterns reflect individual differences in attentional or other general cognitive abilities. The observed patterns of directed connectivity between the hippocampus and parietal cortex may be more closely related to attention and general cognitive processes rather than memory performance. While our study provides valuable insights into the interactions between the hippocampus and parietal cortex during cognitive tasks involving verbal and spatial information processing, further research is necessary to tease out the differential contributions of memory recall and attention. Future studies should incorporate control non-memory tasks to better elucidate the functional significance of these interactions and their specific roles in memory and attentional processes.

### Replication

Finally, a major objective of our study was to probe replicability of findings across verbal and spatial memory domains. Replication of findings is a major problem in neuroscience, particularly in iEEG studies involving multiple distributed brain areas where data sharing is virtually nonexistent and data are difficult to acquire spanning distributed brain regions (Das & Menon, 2022b). To estimate the degree of replicability of our findings across the verbal and spatial memory domains, we employed replication BF (Ly et al., 2019; Verhagen & Wagenmakers, 2014). We found very high replication BF related to the directed connectivity between the hippocampus and the parietal cortex (**Table 4**), emphasizing the robustness of our findings – a challenging proposition in iEEG research. Furthermore, we discovered that most BFs associated with the replication of directed connectivity between the hippocampus and parietal cortex were decisive (BFs > 100), demonstrating consistent results across multiple memory domains and task contexts. It is worth noting that there were some overlaps of subjects across the three experiments that we used (**Figure 1**) and our replication is not a completely independent replication. Future studies with simultaneous dense sampling of electrodes in the hippocampus and the parietal cortex and independent samples of participants across experiments are needed to further examine the replicability of directed connectivity between hippocampus and parietal cortex in human memory processing.

Our findings contribute new insights into the relatively understudied hippocampus-parietal circuits and their role in episodic memory. Remarkably, the patterns of directed connectivity and frequency-specific interactions observed in this study parallel those we previously reported between the hippocampus and the prefrontal cortex (Das & Menon, 2022b). In both cases, we identified asymmetric causal influence during memory encoding and recall periods. This consistency across distinct neural circuits suggests a potentially generalized mechanism for episodic memory formation in the human brain. The robustness of these results, demonstrated through Bayesian analysis, lends further credence to the pivotal role of hippocampus-parietal interactions in episodic memory. Therefore, our current study not only fills a gap in the existing literature but also corroborates and extends our understanding of the complex neural circuit dynamics during memory processing.

### Limitations

Our study has some limitations worth noting. Our study primarily focused on the left hemisphere hippocampus and parietal cortex interactions due to inadequate electrode coverage in the right hippocampus. This limits the understanding of potential hemispheric differences in episodic memory processing. Future studies with denser sampling of electrodes across both hemispheres and multiple verbal and non-verbal memory experiments are needed to separately examine the role of the individual hemispheres during verbal versus non-verbal memory retrieval. While the study included three experiments, there was some overlap of participants across these experiments. A completely independent replication with separate participant groups for each experiment would further strengthen the findings. Due to the limitations of iEEG data acquisition, the study could not analyze interactions between the hippocampus and dorsal parietal subdivisions such as the superior parietal lobule and the intraparietal sulcus. Finally, our conclusions about the causal directed connectivity between the hippocampus and the parietal cortex were assessed using computational methods. To directly establish causal links, future studies should employ causal circuit manipulation techniques such as deep brain stimulation of the hippocampus and parietal cortex, while participants engage in episodic memory tasks.

Despite these limitations, our study provides valuable insights into the electrophysiological dynamics of hippocampal-parietal interactions and their replicability in the human brain.

## Conclusions

Our study advances foundational knowledge of directed connectivity between the hippocampus and the parietal cortex during verbal and spatial memory formation in humans. Utilizing high temporal resolution iEEG recordings from a large cohort of participants and across multiple task domains, we uncovered distinct frequency-specific signaling mechanisms between the hippocampus and the parietal cortex. Our findings not only deepen our understanding of the neural architectures involved in both verbal and spatial memory but also lay groundwork for future studies aimed at exploring memory circuit dysfunctions prominent in psychiatric and neurological disorders. Moreover, our work has implications for future research and potential therapies, particularly those involving targeted stimulation of parietal cortex regions to indirectly modulate hippocampal activity (J. X. Wang et al., 2014). More broadly, our study serves as a reference for deciphering the dynamic electrophysiological interactions that govern complex cognitive functions.

## Acknowledgements

This research was supported by NIH grants NS086085 and EB022907.

